# Differences in syncytia formation by SARS-CoV-2 variants modify host chromatin accessibility and cellular senescence via TP53

**DOI:** 10.1101/2023.08.31.555625

**Authors:** Jonathan D. Lee, Bridget L. Menasche, Maria Mavrikaki, Madison M. Uyemura, Su Min Hong, Nina Kozlova, Jin Wei, Mia M. Alfajaro, Renata B. Filler, Arne Müller, Tanvi Saxena, Ryan R. Posey, Priscilla Cheung, Taru Muranen, Yujing J. Heng, Joao A. Paulo, Craig B. Wilen, Frank J. Slack

## Abstract

COVID-19 remains a significant public health threat due to the ability of SARS-CoV-2 variants to evade the immune system and cause breakthrough infections. Although pathogenic coronaviruses such as SARS-CoV-2 and MERS-CoV lead to severe respiratory infections, how these viruses affect the chromatin proteomic composition upon infection remains largely uncharacterized. Here we used our recently developed integrative DNA And Protein Tagging (iDAPT) methodology to identify changes in host chromatin accessibility states and chromatin proteomic composition upon infection with pathogenic coronaviruses. SARS-CoV-2 infection induces TP53 stabilization on chromatin, which contributes to its host cytopathic effect. We mapped this TP53 stabilization to the SARS-CoV-2 spike and its propensity to form syncytia, a consequence of cell-cell fusion. Differences in SARS-CoV-2 spike variant-induced syncytia formation modify chromatin accessibility, cellular senescence, and inflammatory cytokine release via TP53. Our findings suggest that differences in syncytia formation alter senescence-associated inflammation, which varies among SARS-CoV-2 variants.

## Introduction

Coronavirus disease-2019 (COVID-19) continues to be a threat due to the ongoing evolution of SARS-CoV-2 (severe acute respiratory syndrome coronavirus 2) and concomitant breakthrough infections in vaccinated and previously infected individuals^1–3^. SARS-CoV-2 belongs to the Coronaviridae family of RNA viruses, which includes prior epidemic SARS-CoV-1^4^ and MERS (middle east respiratory syndrome)-CoV^5^ coronaviruses. Despite being in the same family of viruses and causing severe respiratory illness, differences in their genomic sequences and clinical presentations exist^6–8^. Understanding the similarities and differences on host cell states upon infection by different pathogenic coronaviruses and identifying the viral components that bring about these changes may provide insights into the mechanisms underlying their different clinical pathologies.

While nearly all SARS-CoV-2 viral components modulate the host response to infection^9–13^, the SARS-CoV-2 spike in particular has continuously evolved throughout the pandemic, leading to changes not only in immune evasiveness but also in membrane fusogenicity, or the ability for viruses to fuse its membrane with host cell membranes^14–18^. An additional consequence of SARS-CoV-2 spike fusogenicity is the formation of pathological syncytia, produced by cell-cell fusion and observed in some SARS-CoV-2-infected cell cultures and COVID-19 patients^19–24^. Although syncytia formation has been suggested to promote viral replication and transmission^25^, whether differences in syncytia formation via SARS-CoV-2 variants alter host cell responses remains to be determined.

A growing body of evidence suggests that SARS-CoV-2 modifies host cell states upon infection through changes in chromatin states^26–30^. However, how infection via SARS-CoV-2 or other coronaviruses changes the host chromatin proteomic composition remains largely uncharacterized. Using our recently developed integrative DNA And Protein Tagging (iDAPT) chromatin proteomics methodology^31^, we discovered that SARS-CoV-2 infection leads to the stabilization of TP53 protein at open chromatin, which is not observed upon MERS-CoV nor HKU5-SARS-CoV-1-S infection. TP53 contributes to the SARS-CoV-2 host cytopathic effect, an association that is blunted in mutant TP53 cell models. Intracellular expression of the SARS-CoV-2 spike but not the spikes of other human coronaviruses recapitulates the TP53 protein stabilization observed upon infection. We found that TP53 stabilization is a consequence of syncytia formation via the SARS-CoV-2 spike: TP53 protein levels are increased with increasingly fusogenic SARS-CoV-2 spike sequences and diminished by inhibition of syncytia formation. Differences in spike-induced TP53 stabilization modify chromatin accessibility states, activation of cellular senescence, the senescence-associated secretory phenotype (SASP), and the release of inflammatory cytokines. Finally, among a panel of SARS-CoV-2 spike variants, differences in syncytia formation correlate with differences in TP53 stabilization. Our findings suggest that virus-induced syncytia formation may contribute to inflammation via the activation of TP53-dependent cellular senescence and SASP-induced inflammation, which may continue to vary with future SARS-CoV-2 variants and nascent coronaviruses.

## Results

### TP53 protein is stabilized upon SARS-CoV-2 infection and contributes to the SARS-CoV-2 host cytopathic effect

To examine differences in coronaviral infection on host cell states, we used the VeroE6 cell line model, which is permissive to SARS-CoV-2 and MERS-CoV infection^32^ and largely recapitulates the host transcriptomic features of SARS-CoV-2 infection observed with other cell-based models (**Fig. S1A-B**)^33,34^. In addition to viral isolates of SARS-CoV-2 and MERS-CoV, we used HKU5-SARS-CoV-1-S as a model of SARS-CoV-1, which encodes the SARS-CoV-1 spike protein^35^. As our primary objective was to elucidate the key regulators of host chromatin state changes upon infection, we used the recently developed iDAPT-MS (integrative DNA and Protein Tagging with mass spectrometry)^31^ in tandem with the Assay for Transposase-Accessible Chromatin (ATAC)-seq^36^ to identify changes in chromatin proteomic composition and corresponding chromatin accessibility states (**Fig. 1A**). We found that SARS-CoV-2 infection induces pervasive changes across the chromatin accessibility landscape (**Fig. 1B**). Concordant with the regulatory capacity of chromatin accessibility on transcription^37,38^, genes with increased accessibility at their transcriptional start sites (TSS’s) are associated with increased gene expression; however, genes with decreased TSS accessibility do not broadly exhibit decreased gene expression (**Fig. S1C**). When comparing the locus-specific chromatin accessibility changes induced by the three coronaviruses, we observed distinct separation of mock-infected cells from virus-infected cells along the first principal component (**Fig. 1C, S1D-E**). The three viruses differ from each other along the second principal component, with the greatest separation between SARS-CoV-2 and MERS-CoV infection; SARS-CoV-2 and HKU5-SARS-CoV-1-S ATAC-seq profiles further separate along components 3 and 4 (**Fig. 1C, S1F**). Transcription factor motif analysis yields a similar distinction between the three viral infections (**Fig. S1G**). Thus, while the three coronaviruses broadly induce similar changes to chromatin accessibility profiles upon infection relative to mock infection, we also observed coronavirus-specific effects on host chromatin accessibility profiles upon infection.

**Figure 1.**
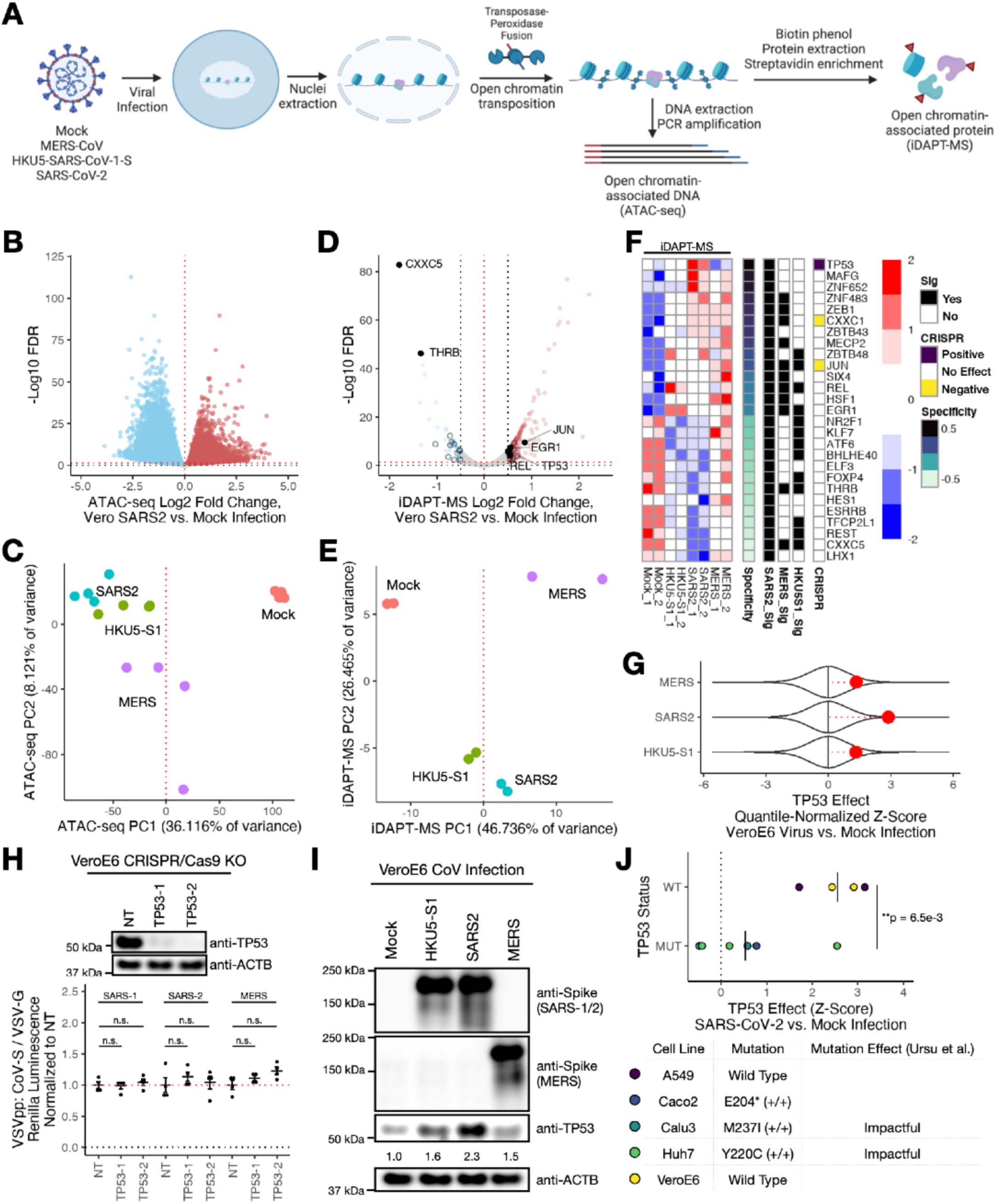
TP53 protein is stabilized upon SARS-CoV-2 infection and contributes to the SARS-CoV-2 host cytopathic effect. A. Experimental design and schematic of the iDAPT approach (made with Biorender). B. ATAC-seq volcano plot demonstrating the changes in accessible chromatin loci upon SARS-CoV-2 infection versus mock infection in VeroE6 cells 48 hr post-infection. FDR, false discovery rate. C. Principal component analysis of ATAC-seq profiles upon infection with SARS-CoV-2, HKU5-SARS-CoV-1-S, or MERS-CoV and mock infection. PC, principal component. D. iDAPT-MS volcano plot of SARS-CoV-2 infection (MOI 0.5) versus mock infection in VeroE6 cells 48 hr post-infection. Black circles, significant transcription factors (FDR < 0.05 and |log2 Fold Change| > 0.5). E. Principal component analysis of iDAPT-MS profiles upon infection with SARS-CoV-2, HKU5-SARS-CoV-1-S, or MERS-CoV and mock infection. F. Heatmap of relative protein abundance levels of differentially abundant (FDR < 0.05 and |log2 Fold Change| > 0.5) transcription factors from SARS-CoV-2 vs. mock infection iDAPT-MS analyses. Specificity is defined as the Pearson correlation between iDAPT-MS protein abundance and SARS-CoV-2 infection status. Sig, significant differential chromatin abundance by iDAPT-MS (FDR < 0.05 and |log2 Fold Change| > 0.5) upon infection relative to mock-infected cells. CRISPR, VeroE6 SARS-CoV-2 CRISPR/Cas9 screening results from Wei et al^32^. Positive, Z-score > 2. Negative, Z-score < -2. G. TP53 sgRNA enrichment from CRISPR/Cas9 pooled screening in VeroE6 cells for the three coronaviruses assessed versus mock infection (Wei et al^32^). Z-scores are quantile-normalized for improved comparison between distributions. H. Top, Western blotting analysis of the indicated proteins after CRISPR/Cas9-based targeting of two TP53-targeting sgRNAs (TP53-1 and TP53-2) or negative control (NT, nontargeting). Bottom, the corresponding Vero E6 cells were infected with VSV pseudotyped viral particles (VSVpp) expressing VSV-G or different coronaviral spike proteins. Luciferase relative to the VSV-G-expressing VSVpp control was measured 24 h after infection. The mean ± s.e.m. are shown. The Welch two-sample t-test with Holm p-value correction was used to assess statistical significance relative to negative control: n.s., not significant. I. Western blotting analysis of the indicated proteins after infection with the corresponding coronaviruses (MOI 0.1, 24 h). Relative ratios of TP53/ACTB (normalized to mock) are annotated below TP53. J. Top, TP53 sgRNA enrichment across published CRISPR/Cas9 genetic screens stratified by TP53 mutational status of the parental cell line (Rebendenne et al^54^). The mean Z-score of each group is demarcated by a horizontal line. p-value, Welch two-sample t-test. Bottom, TP53 mutational status associated with each parental cell line and corresponding impact of TP53 point mutations on its transcriptional activity as determined in Ursu et al.^62^.

Given the widespread and differential changes to chromatin accessibility upon coronavirus infection, we sought to define the compositional changes to the chromatin proteome by iDAPT-MS, which may nominate key mediators of these chromatin accessibility differences. Consistent with chromatin accessibility and transcriptomic shifts observed upon SARS-CoV-2 infection^27,33,34,39^, we identified numerous changes in host chromatin proteomic composition, including 14 transcription factors with increased chromatin abundance and 13 with decreased abundance relative to mock-infected cells (**Fig. 1D, S2A-C).**

Comparison between iDAPT-MS profiles of coronavirus-infected and mock-infected cells shows separation of virus infection from mock along the first principal component (**Fig. 1E**). SARS-CoV-2 and HKU5-SARS-CoV-1-S profiles segregate from MERS-CoV profiles along the second principal component, and SARS-CoV-2 and HKU5-SARS-CoV-1-S profiles separate along the third principal component, reflecting the differences between coronavirus infections observed by ATAC-seq (**Fig. 1C, 1E, S2D**). Among differentially abundant transcription factors common to all three coronavirus infections, EGR1, REL, and JUN are significantly increased, whereas THRB and CXXC5 are decreased on chromatin (**Fig. 1D, S2E-F**). These transcription factors have been implicated in the host cell response to viral infection and cellular stress. For instance, EGR1 and JUN regulate cell proliferation, immune responses, and cell death and may be activated upon viral infection including many coronaviruses^40–42^, whereas REL, a member of the NF-kappaB family of transcription factors, potentiates immune and inflammatory responses including upon SARS-CoV-2 infection^27,43^. CXXC5 modulates cellular metabolism, proliferation, and death; moreover, it may also modify interferon responses upon viral infections via DNA methylation^44,45^. Finally, THRB mediates the transcriptional activity of thyroid hormones and has also been reported to modulate innate and adaptive immunity^46^.

Next, we sought to identify transcription factors that change in chromatin abundance specifically upon SARS-CoV-2 infection. iDAPT-MS reveals SARS-CoV-2-specific increases of 4 transcription factors (TP53, MAFG, ZNF652, ZBTB43) and decreases of 4 transcription factors (LHX1, ESRRB, HES1, ELF3) relative to mock-infected cells (**Fig. 1F**). Of these, not only is TP53 the transcription factor most specifically enriched by iDAPT-MS upon SARS-CoV-2 infection, but it is also the only transcription factor that substantially contributes to the SARS-CoV-2 cytopathic effect in VeroE6 cells, measured by CRISPR/Cas9 sgRNA dropout upon infection^32^ (**Fig. 1F**). Moreover, TP53 effects on the cytopathic effect are diminished in both MERS-CoV and HKU5-SARS-CoV-1-S infections (**Fig. 1G**). TP53 loss does not affect the entry of vesicular stomatitis virus (VSV) pseudoviral particles (VSVpp) displaying the SARS-CoV-1, SARS-CoV-2, or MERS-CoV coronavirus spike glycoproteins, in support of its effects on the cytopathic effect downstream of viral entry (**Fig. 1H**)^47,48^. Finally, TP53 protein levels are increased upon SARS-CoV-2 infection and less so upon HKU5-SARS-CoV-1-S and MERS-CoV infections, consistent with its regulation via protein stability^49^ (**Fig. 1I**). Altogether, our analyses indicate that TP53 protein stabilization and activity are SARS-CoV-2-specific consequences of infection in VeroE6 cells.

That TP53 is uniquely stabilized by SARS-CoV-2 among coronavirus infections was unexpected. On the one hand, TP53 is a general stress factor often induced by viral infections, and as such we might expect TP53 up in all three viral infection conditions^50–52^. On the other hand, coronavirus-encoded nsp3 has been suggested to antagonize TP53 stabilization^53^. Given its surprisingly unique enrichment upon SARS-CoV-2 infection combined with its SARS-CoV-2-specific effects on the host cytopathic effect, we sought to further investigate the role of TP53 on the host cytopathic effect.

We assessed whether SARS-CoV-2-associated TP53 effects are specific to VeroE6 cells or broadly consequential in other cell contexts. First, we observed positive enrichment of a TP53 target gene signature among transcriptomic changes observed upon SARS-CoV-2 infection in VeroE6, A549ACE2, and COVID-19 patient ATII pulmonary cells^33,34,39^ (**Fig. S2G**). Second, across CRISPR/Cas9 SARS-CoV-2 screening datasets^32,54–60^, TP53 has a consistent effect in promoting the host cytopathic effect (**Fig. S2H**). Third, because TP53 is often mutated or deleted in cell line models, we stratified these CRISPR/Cas9 screening datasets by TP53 mutational status, which may fully or partially inactivate its transcriptional activity^61,62^. Indeed, TP53 consistently promotes the cytopathic effect upon SARS-CoV-2 infection in wild-type TP53 cell lines (VeroE6 and A549) but not in TP53-deleted (Caco2) or point mutation-inactivating (Calu3, Huh7) cell lines (**Fig. 1J**). Altogether, our analyses indicate that TP53 contributes to the host cell state changes and cytopathic effects induced by SARS-CoV-2 infection.

Finally, we expanded our functional iDAPT-MS analyses beyond transcription factors. Among those genes which modulate the SARS-CoV-2 host cytopathic effect in VeroE6 cells and are significant by iDAPT-MS, we found consistent effects of both ELL and PRPF39 on the SARS-CoV-2 cytopathic effect in other cellular contexts (**Fig. S2H**). ELL, an RNA polymerase II elongation factor implicated in cellular stress responses^63,64^, increases by iDAPT-MS and antagonizes the host cytopathic effect. On the other hand, PRPF39, which goes down by iDAPT-MS while also antagonizing the cytopathic effect, may realize its functions through its transcriptional splicing activity^65^. However, these effects are not specific to SARS-CoV-2 infection by iDAPT-MS. On the other hand, we found SGO1 as a SARS-CoV-2-specific protein by iDAPT-MS, but its effect on the cytopathic effect was not consistent in other screening datasets. Given the SARS-CoV-2 specificity and consistency of TP53 activity across multiple datasets not observed with other proteins, we proceeded with TP53 analysis in SARS-CoV-2 infection.

### The SARS-CoV-2 spike protein is a key determinant of host cell chromatin accessibility changes and TP53 stabilization

In light of our findings, we hypothesized that differences in coronavirus protein sequences contribute to the observed differences in chromatin accessibility profiles and TP53 stabilization. To test this, we first profiled chromatin accessibility profiles of VeroE6 cells upon expression of each of the 29 SARS-CoV-2 viral proteins^13,66^ (**Fig. 2A, S3A-B**). Among these proteins, SARS-CoV-2 nsp1 and spike proteins promote widespread changes in chromatin accessibility, and these changes significantly and positively correlate with the changes observed upon SARS-CoV-2 viral infection (**Fig. 2A-C**). However, the chromatin accessibility changes induced by nsp1 and spike differ from each other, with minimal overlap between the two profiles; critically, only spike protein expression leads to TP53 protein stabilization (**Fig. 2D-E**). Direct comparison with SARS-CoV-1 and MERS-CoV spike proteins reveals that TP53 stabilization occurs only with the SARS-CoV-2 spike protein (**Fig. 2F**). We expanded this analysis to the four other human coronaviruses, 229E, HKU1, NL63, and OC43^67^: none of these spike sequences are sufficient to stabilize TP53 to the level of the SARS-CoV-2 spike (**Fig. 2F**). Thus, the SARS-CoV-2 spike versus other human coronavirus spike proteins uniquely contributes to host cell TP53 stabilization.

**Figure 2.**
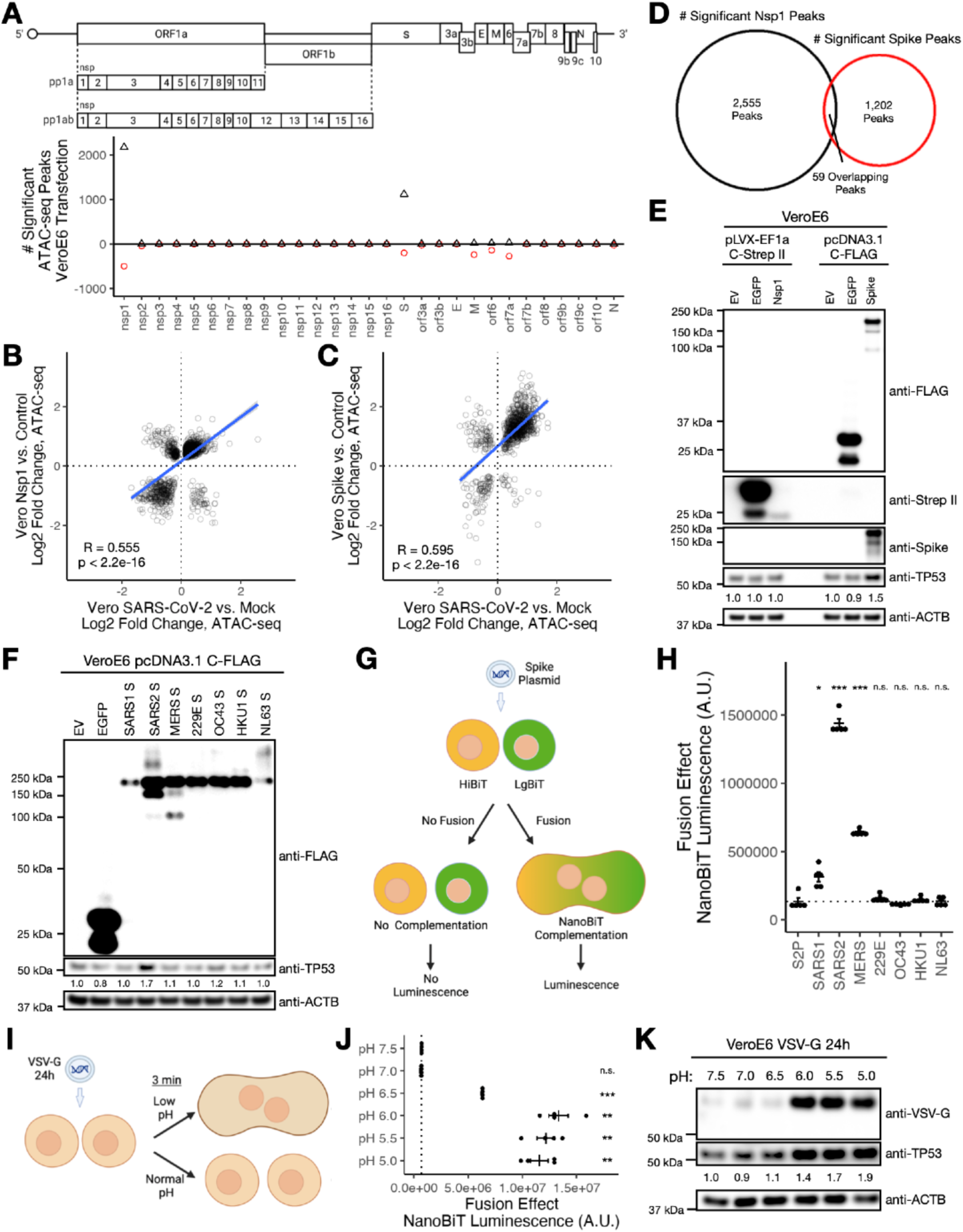
The SARS-CoV-2 spike protein is a key determinant of host cell chromatin accessibility changes and TP53 stabilization. A. Top, schematic of the SARS-CoV-2 viral genome with encoded proteins. Bottom, number of significant (FDR < 0.05) ATAC-seq peaks upon transfection of plasmid-encoded viral proteins relative to empty vector and EGFP (48 hr) in VeroE6 cells. Black triangles, peaks with increasing accessibility. Red circles, peaks with decreasing accessibility. B. Comparison of thresholded (FDR < 0.05) differential ATAC-seq peaks between SARS-CoV-2 vs. mock infection (24 hr, MOI 0.5) and plasmid-encoded transfection of SARS-CoV-2 nsp1 vs. controls in VeroE6 cells (48 hr). Blue line, best-fit trendline. R, Pearson correlation. p-value, Pearson correlation p-value. C. Comparison of thresholded (FDR < 0.05) differential ATAC-seq peaks between SARS-CoV-2 vs. mock infection (24 hr, MOI 0.5) and plasmid-encoded transfection of SARS-CoV-2 spike vs. controls in VeroE6 cells (48 hr). Blue line, best-fit trendline. R, Pearson correlation. p-value, Pearson correlation p-value. D. Venn diagram analysis of differential ATAC-seq peaks (FDR < 0.05, positive and negative combined) upon nsp1 or spike expression relative to controls. E. Western blotting analysis of VeroE6 cells 48 hr after transfection with plasmids encoding the indicated proteins. EV, empty vector. Relative ratios of TP53/ACTB (normalized to the corresponding EV) are annotated below TP53. F. Western blotting analysis of VeroE6 cells 48 hr after transfection with plasmids encoding the indicated FLAG-tagged proteins. EV, empty vector. Relative ratios of TP53/ACTB (normalized to EV) are annotated below TP53. G. Schematic of cell-cell fusion assay. A 1:1 mixture of VeroE6 cells stably transduced with either pLenti-HiBiT-Bsr or pLenti-LgBiT-P2A-Bsr is transfected with corresponding plasmids. After 48 hr, the cell-cell fusion effect is measured by NanoBiT luciferase complementation. Made with Biorender. H. Cell-cell fusion analysis of VeroE6 cells transfected with the corresponding spike proteins. S2P, “2P” mutant of the spike which is incapable of membrane fusion. The Welch two-sample t-test with Holm p-value correction was used to assess statistical significance relative to the S2P mutant as negative control: ***, p < 0.001; *, p < 0.05; n.s., not significant. I. Schematic of VSV-G acidification assay. Cells were transfected with plasmid encoding VSV-G. 24 h later, cells were treated with a 3 min pulse of DMEM buffered with 10 mM morpholineethanesulfonic acid (MES) at different pH levels. Cells were analyzed 24 h after acidification. J. Cell-cell fusion analysis of VeroE6 cells transfected with VSV-G plasmid and treated with buffers at different pH levels. The Welch two-sample t-test with Holm p-value correction was used to assess statistical significance relative to pH 7.5 as control: ***, p < 0.001; **, p < 0.01; n.s., not significant. K. Western blotting analysis of VeroE6 cells transfected with VSV-G plasmid and treated with buffers at different pH levels. Relative ratios of TP53/ACTB (normalized to pH 7.5) are annotated below TP53.

A characteristic of the SARS-CoV-2 spike is its increased propensity to fuse cells together and form multinucleated syncytia^19–24^. Indeed, by a NanoBiT luciferase complementation assay, SARS-CoV-2 spike promotes substantially increased cell-cell fusion relative to other human coronavirus spike proteins, whereas nsp1 does not promote cell-cell fusion (**Fig. 2G-H, S3C-D**). To assess if TP53 stabilization is a SARS-CoV-2 spike-specific effect or a consequence of virus-induced membrane fusion in VeroE6 cells, we tested whether the acidification of the VSV glycoprotein (VSV-G), which induces plasma membrane fusion, would also stabilize TP53 (**Fig. 2I**)^68,69^. We found that VeroE6 cells exhibit increased TP53 protein levels in a pH-dependent manner (**Fig. 2J-K, S3E**). Of note, despite cells being subject to the same amount of transfected VSV-G plasmid, VSV-G protein levels upon acidification are substantially increased, possibly reflecting increased protein stability at the plasma membrane versus its typical lysosomal degradation (**Fig. 2K**)^70,71^. Altogether, our findings are consistent with a relationship between virus-induced cell-cell fusion and TP53 protein stabilization, which varies among pathogenic coronaviruses due to differences in their spike proteins.

### Increasingly fusogenic SARS-CoV-2 variants promote increased TP53 protein stabilization

Since the start of the COVID-19 pandemic, mutations in the SARS-CoV-2 spike protein have emerged that not only facilitate evasion of host immune responses but also modulate cell-cell fusogenicity^14–16,19,72,73^. Due to the increased fusogenicity of the alpha/B.1.1.7 and delta/B.1.617.2 SARS-CoV-2 spike sequences, we assessed whether the increases in syncytia formation leads to increased TP53 stabilization (**Fig. 3A**). Indeed, TP53 protein levels are increased upon alpha and delta spike expression relative to ancestral spike in VeroE6 cells (**Fig. 3B**). In A549ACE2 cells, which are susceptible to spike-induced cell-cell fusion, we also detected ancestral spike-dependent TP53 stabilization that is further increased by the delta spike (**Fig. 3C-D**).

**Figure 3.**
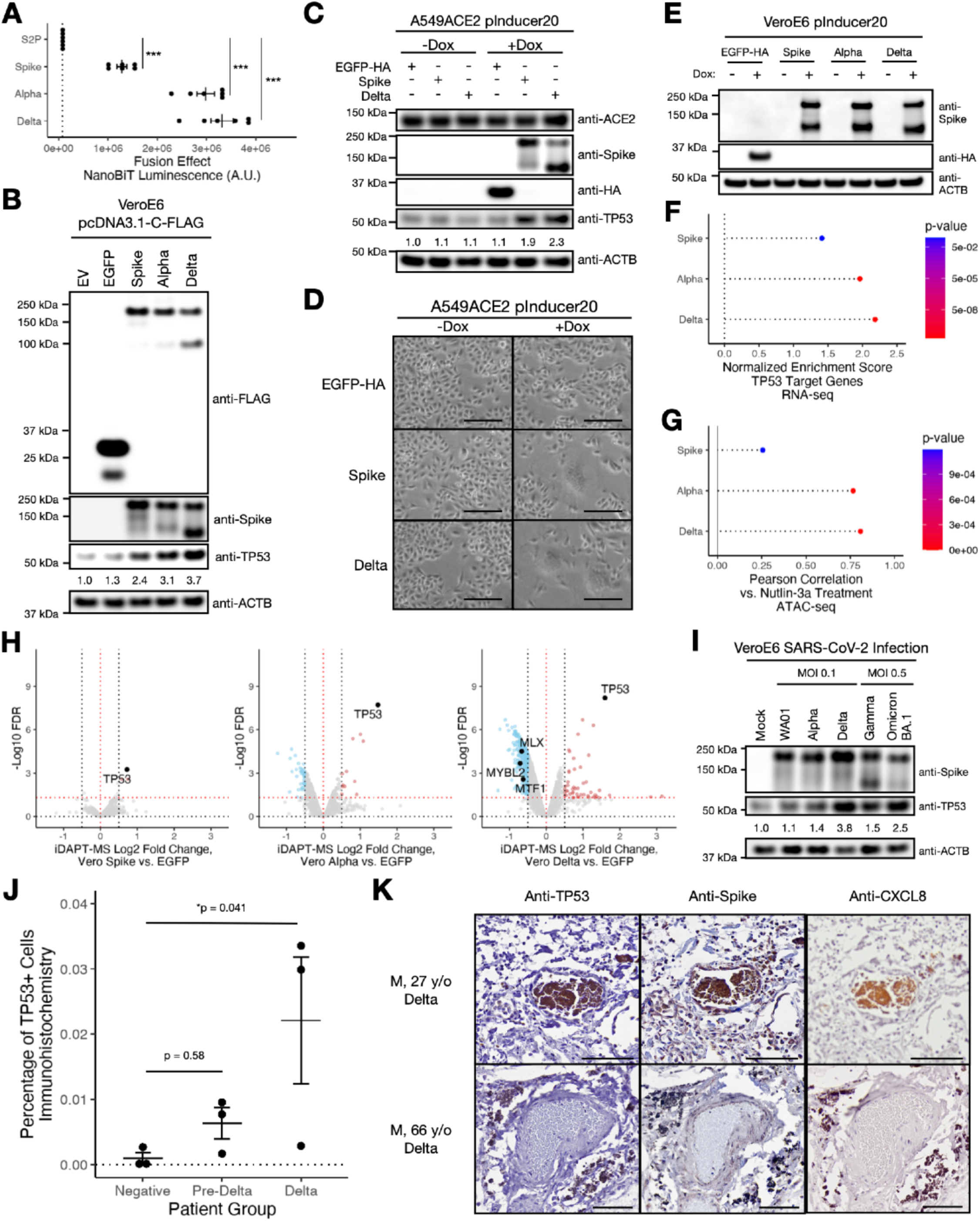
Increasingly fusogenic SARS-CoV-2 variants promote increased TP53 protein stabilization. A. Cell-cell fusion analysis of VeroE6 cells transfected with the corresponding spike proteins. S2P, “2P” mutant of the spike. The Welch two-sample t-test with Holm p-value correction was used to assess statistical significance relative to the S2P mutant as negative control: ***, p < 0.001. B. Western blotting analysis of VeroE6 cells 48 hr after transfection with plasmids encoding the indicated FLAG-tagged proteins. EV, empty vector. Relative ratios of TP53/ACTB (normalized to EV) are annotated below TP53. C. Western blotting analysis of A549ACE2 pInducer20 cells transduced with the indicated gene constructs and treated with or without 1 μg/mL doxycycline (Dox) for 48 hr. Relative ratios of TP53/ACTB (normalized to EGFP -Dox) are annotated below TP53. D. Representative brightfield images of A549ACE2 pInducer20 cells transduced with the indicated gene constructs and treated with or without 1 μg/mL Dox for 48 hr. Scale bar, 200 μm. E. Western blotting analysis of the indicated proteins 48 hr after 1 μg/mL Dox treatment of VeroE6 pInducer20 cells transduced with the indicated gene constructs. F. RNA-seq analysis of TP53 target gene upon ancestral, alpha, or delta spike expression versus EGFP expression. p-value, gene set enrichment analysis. Also see **Fig. S4B, S4G**. G. Pearson correlation of ATAC-seq profiles upon ancestral, alpha, or delta spike expression versus EGFP expression compared to treatment with or without 1 μM nutlin-3a for 24 hr. p-value, Pearson correlation. Also see **Fig. S4C-F**. H. iDAPT-MS volcano plots of ancestral, alpha, or delta spike expression versus EGFP expression in VeroE6 pInducer20 cells 48 hr after 1 μg/mL doxycycline induction. Black points, significant transcription factors (FDR < 0.05 and |log2 Fold Change| > 0.5). I. Western blotting analysis of the indicated proteins after infection with the corresponding SARS-CoV-2 viral variants and MOIs (24 h). Relative ratios of TP53/ACTB (normalized to mock) are annotated below TP53. J. Percentage of TP53-positive cells in COVID-19 patients with pre-Delta or Delta SARS-CoV-2 variants versus uninfected (negative) patient lung biopsies. Immunohistochemistry quantification was performed using QuPath. p-value, linear regression coefficient. K. Representative immunohistochemistry images of TP53 protein, SARS-CoV-2 spike, and CXCL8/IL8 protein epitopes in the lungs of Delta variant-infected COVID-19 patients. Scale bar, 100 μm.

Next, we sought to determine the effects of SARS-CoV-2 spike expression on host cell states and how they may differ with differences in variant spike-induced syncytia formation. Doxycycline-induced spike expression in VeroE6 cells after 48 hours leads to similar levels of total spike protein expression for ancestral, alpha, and delta sequences, with increased syncytia formation observed with alpha and delta variants relative to the ancestral spike sequence^19,73^ (**Fig. 3E, S4A**). To assess how the SARS-CoV-2 spike affect chromatin states, we performed “Multilevel Chromatin Analysis”, which includes tandem transcriptome (RNA-seq), chromatin accessibility (ATAC-seq), and chromatin proteome (iDAPT-MS) analyses. At all three levels of chromatin analysis, ancestral spike yields the weakest effects relative to EGFP expression, whereas alpha and delta spike sequences promote the most dramatic changes (**Fig. 3F-H, S4B-C**). We found these increases in effect sizes to correspond to increases in TP53 target gene expression (RNA-seq), nutlin-3a-associated chromatin accessibility signatures (ATAC-seq), and TP53 chromatin protein enrichment (iDAPT-MS) (**Fig. 3F-H, S4B-G, S5A-B**). Surprisingly, TP53 is the only sequence-specific transcription factor with increased abundance on open chromatin in all three spike-expression settings (**Fig. 3H**). We further validated the increased TP53 abundance in the nucleus upon expression of different spike variants by subcellular fractionation (**Fig. S5C**). Thus, Multilevel Chromatin Analysis nominates TP53 as a key contributor to SARS-CoV-2 spike-driven host cell state changes, which varies with differences in spike sequence and fusogenicity.

We assessed whether these spike-mediated differences in TP53 stabilization are observed upon bona fide SARS-CoV-2 viral infection. Indeed, we found that infection with a delta isolate of the virus leads to much stronger stabilization in infected VeroE6 cells (**Fig. 3I**). Furthermore, lung autopsy samples of COVID-19 patients infected with the delta variant show an increased frequency of cells positively staining for TP53, signal which is proximal to SARS-CoV-2 spike protein epitope signal (**Fig. 3J-K**). Thus, our findings are in support of TP53 stabilization as a consequence not only of fusogenic spike expression but also of SARS-CoV-2 viral infection, which varies with the fusogenicity of the viral spike sequence.

### Inhibition of SARS-CoV-2 spike-induced syncytia formation attenuates TP53 stabilization

Although we have thus far observed an association between the magnitude of TP53 activation and cell-cell fusion, driven by differences in the fusogenicity of the SARS-CoV-2 spike protein, we sought to test this mechanism formally. First, we found that despite productive spike/ACE2 interaction and spike-dependent viral entry as denoted by Renilla luciferase expression, VSVpp particles expressing the SARS-CoV-2 spike glycoprotein do not induce stabilization of host TP53 in A549ACE2 cells, even when the spike sequence is replaced with alpha or delta variant sequences (**Fig. 4A**). We also found that spike expression in A549 parental cells, which are negative for ACE2 and do not permit cell-cell fusion, does not increase TP53 protein levels higher than the expression of EGFP (**Fig. S5D-E**). On the other hand, 1:1 mixtures of these spike-expressing A549 cells with A549ACE2-EGFP cells, which alone do not stabilize TP53 nor promote cell-cell fusion, promote substantial TP53 protein stabilization and syncytia formation (**Fig. 3C, 4B, S5D**). Consistent with cell-cell interaction promoting TP53 activation, a neutralizing antibody against the delta spike that blocks cell-cell fusion also diminishes TP53 levels induced by delta spike-expressing A549ACE2 cells (**Fig. 4C**).

**Figure 4.**
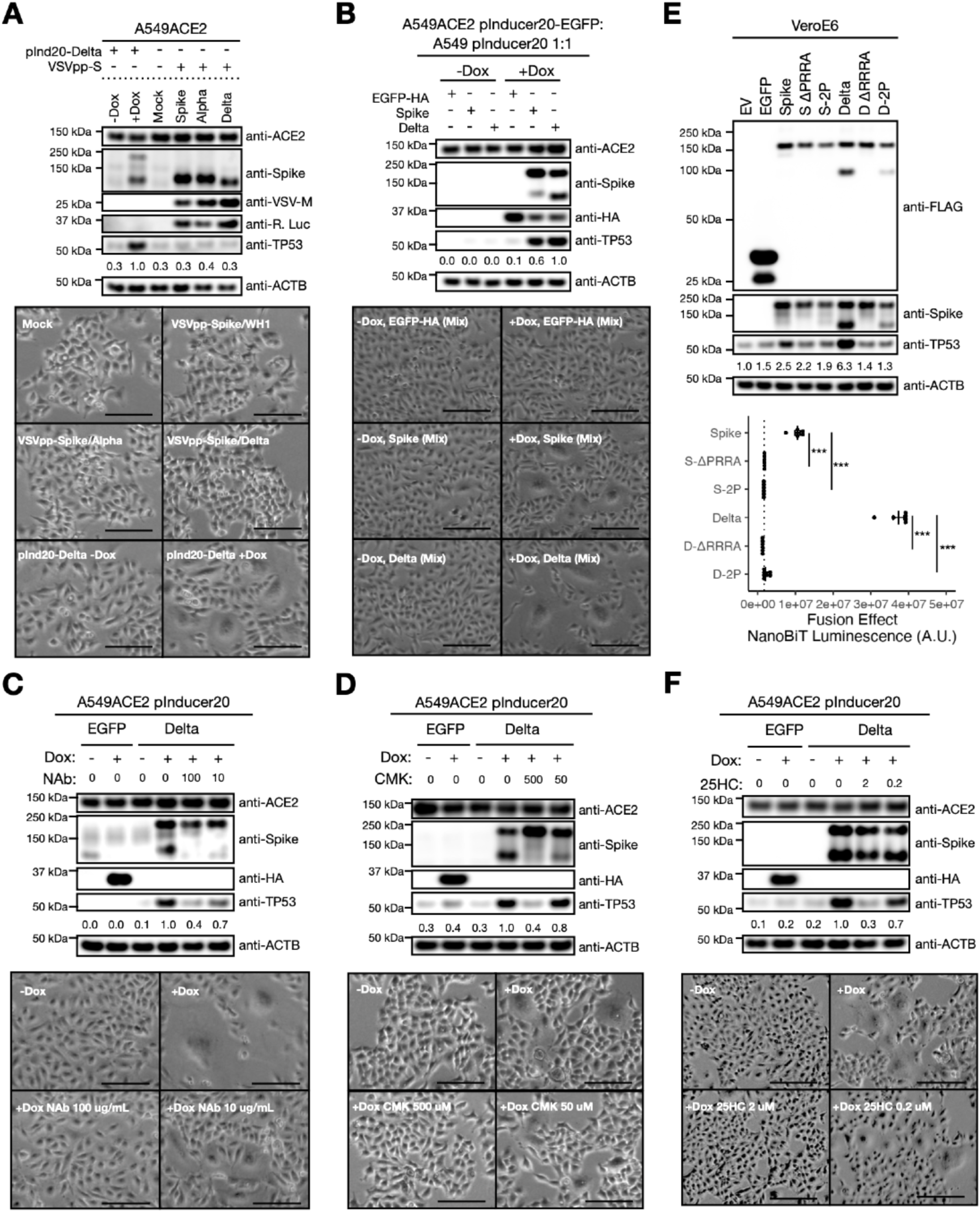
Inhibition of SARS-CoV-2 spike-induced syncytia formation attenuates TP53 stabilization. A. Top, western blotting analysis of A549ACE2 cells infected with VSVpp-SARS-CoV-2-S (VSVpp-S) displaying the indicated SARS-CoV-2 spike sequences for 24 hr. A549ACE2 pInducer20 cells transduced with the Delta spike sequence (pInd20-Delta) and treated with or without 1 μg/mL doxycycline (Dox) for 24 hr are shown for comparison. Relative ratios of TP53/ACTB (normalized to Delta +Dox) are annotated below TP53. Bottom, representative brightfield images of A549ACE2 cells infected with the corresponding VSVpp-SARS-CoV-2-S viruses for 24 hr. Scale bar, 200 μm. B. Top, western blotting analysis of 1:1 mixtures of A549 and A549ACE2 pInducer20 cells transduced with the indicated gene constructs and treated with or without 1 μg/mL Dox for 48 hr. Relative ratios of TP53/ACTB (normalized to Delta +Dox) are annotated below TP53. Bottom, representative brightfield images of A549/A549ACE2 pInducer20 cell mixtures as above treated with or without 1 μg/mL Dox for 48 hr. Scale bar, 200 μm. C. Top, western blotting analysis of A549ACE2 pInducer20 cells transduced with the indicated gene constructs and treated with or without 1 μg/mL Dox and with the corresponding amount of neutralizing antibody (NAb, μg/mL, Sino Biological #40592-R001) for 24 hr. Relative ratios of TP53/ACTB (normalized to Delta +Dox) are annotated below TP53. Bottom, representative brightfield images of A549ACE2 pInducer20-Delta spike cells treated with or without 1 μg/mL Dox and with the corresponding amount of neutralizing antibody (μg/mL) for 24 hr. Scale bar, 200 μm. D. Top, western blotting analysis of A549ACE2 pInducer20 cells transduced with the indicated gene constructs and treated with or without 1 μg/mL Dox and with the corresponding amount of CMK (μM) for 24 hr. Relative ratios of TP53/ACTB (normalized to Delta +Dox) are annotated below TP53. Bottom, representative brightfield images of A549ACE2 pInducer20-Delta spike cells treated with or without 1 μg/mL Dox and with the corresponding amount of CMK (μM) for 24 hr. Scale bar, 200 μm. E. Top, western blotting analysis of VeroE6 cells 48 hr after transfection with plasmids encoding the indicated FLAG-tagged proteins. Relative ratios of TP53/ACTB (normalized to EV) are annotated below TP53. Bottom, cell-cell fusion analysis of VeroE6 cells transfected with the corresponding spike proteins. S-2P/D-2P, “2P” mutant of the spike which is incapable of membrane fusion. The Welch two-sample t-test with Holm p-value correction was used to assess statistical significance relative to the S-2P mutant as negative control: ***, p < 0.001. F. Top, western blotting analysis of A549ACE2 pInducer20 cells transduced with the indicated gene constructs and treated with or without 1 μg/mL Dox and with the corresponding amount of 25-hydroxycholesterol (μM) for 24 hr. Relative ratios of TP53/ACTB (normalized to Delta +Dox) are annotated below TP53. Bottom, representative brightfield images of A549ACE2 pInducer20-Delta spike cells treated with or without 1 μg/mL Dox and with the corresponding amount of 25-hydroxycholesterol (μM) for 24 hr. Scale bar, 200 μm.

Given the spike protein’s processing by host cell proteases, we tested protease inhibitors for their effects on host TP53 activation. Treatment with high concentrations of CMK, a FURIN protease inhibitor that prevents S1/S2 spike cleavage at the polybasic cleavage site^74^, abolishes TP53 stabilization and cell-cell fusion (**Fig. 4D**). Indeed, deletion of the polybasic cleavage sites in both ancestral (PRRA) and delta (RRRA) spike^72,75–77^ sequences leads to loss of TP53 stabilization and membrane fusion (**Fig. 4E**). On the other hand, the cathepsin L inhibitor E64d^78^ does not affect TP53 stabilization at the concentrations tested (**Fig. S5F**). We found that mutating the spike amino acid residues L986 and V987 to two proline residues (“S-2P”/”D-2P” mutants), which prevents the spike protein from accessing the postfusion conformational state^79,80^, abrogates TP53 stabilization and cell-cell fusion (**Fig. 4E**). Furthermore, 25-hydroxycholesterol and imatinib, both of which perturb cell-cell fusion via host cell mechanisms, inhibit delta spike-induced TP53 activation (**Fig. 4F, S5G**)^81–85^. Our findings indicate that cell-cell fusion induced by the SARS-CoV-2 spike are required for TP53 stabilization.

### Differences in spike-induced TP53 stabilization modify the activation of host chromatin accessibility, senescence, and inflammatory cytokine release

Thus far, we have found that TP53 protein is stabilized due to spike-induced cell-cell fusion and that the magnitude of its stabilization varies with the degree of fusion. We next sought to determine how TP53 contributes to spike-dependent cellular phenotypes. Although stabilization of TP53 is a consequence of syncytia formation (**Fig. 4**) and its genetic loss modulates the host cytopathic effect (**Fig. 1J**), TP53 loss itself does not block syncytia formation (**Fig. 5A-B**).

**Figure 5.**
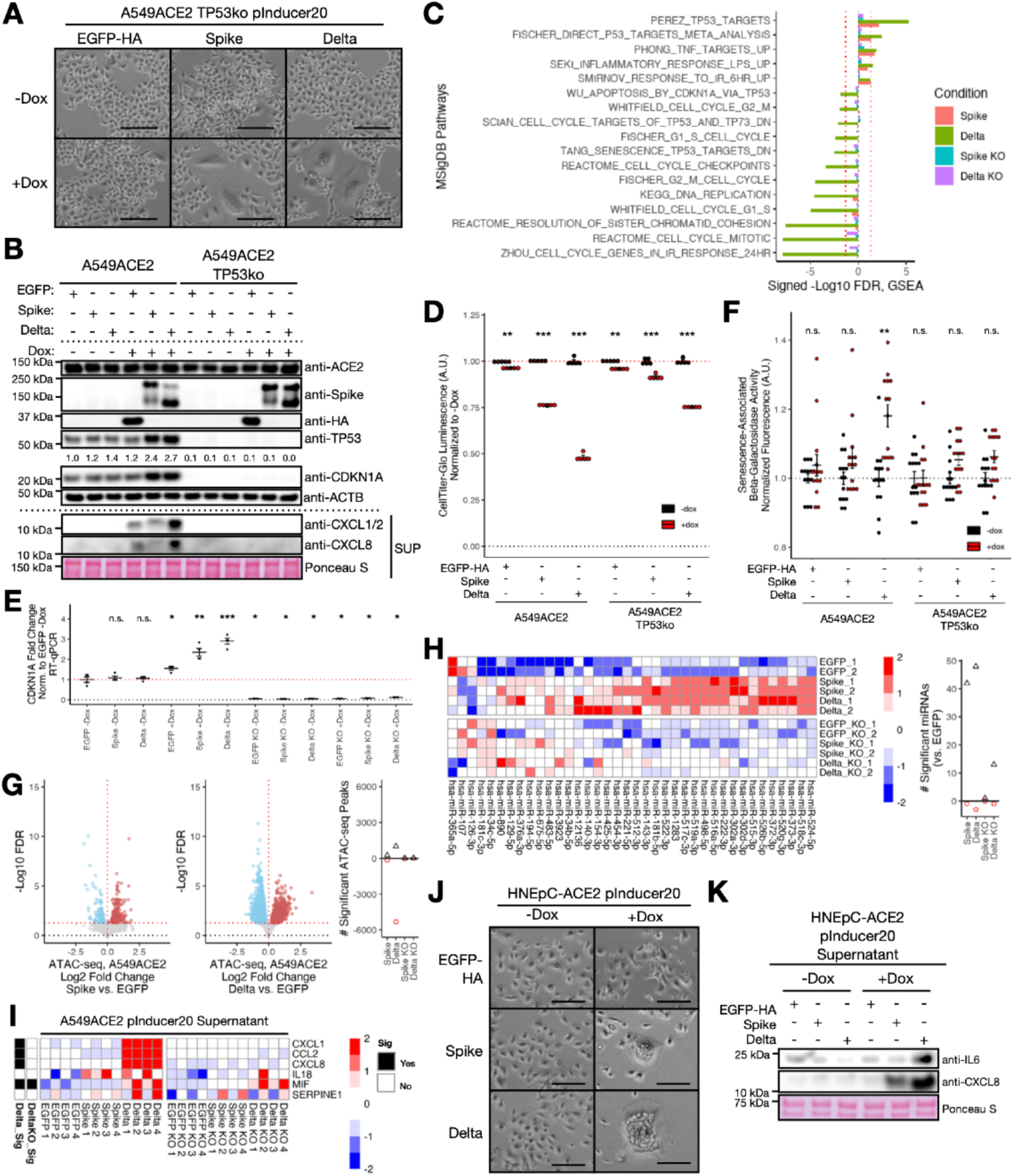
Differences in spike-induced TP53 stabilization modify the activation of host chromatin accessibility, senescence, and inflammatory cytokine release. A. Representative brightfield images of A549ACE2 TP53ko pInducer20 cells transduced with the indicated gene constructs and treated with or without 1 μg/mL Dox for 48 hr. Scale bar, 200 μm. B. Top, Western blotting analysis of whole cell lysates of A549ACE2 or A549ACE2 TP53ko pInducer20 cells transduced with the indicated gene constructs and treated with or without 1 μg/mL Dox for 48 hr. Relative ratios of TP53/ACTB (normalized to EGFP -Dox) are annotated below TP53. Bottom, Western blotting analysis of supernatants (SUP) of the corresponding cell constructs and treatments. C. MSigDB pathway enrichment analysis of differential RNA-seq profiles upon Dox-induced expression (48 hr, 1 μg/mL) of SARS-CoV-2 ancestral or delta spike relative to expression of EGFP in either A549ACE2 or A549ACE2 TP53ko pInducer20 cells. D. Cell viability as assessed by CellTiter-Glo in the indicated A549ACE2 or A549ACE2 TP53ko pInducer20 cells treated with or without 1 μg/mL Dox for 72 hr (mean ± s.e.m.). The Welch two-sample t-test with Holm p-value correction was used to assess statistical significance: ***, p < 0.001; **, p < 0.01. E. RNA levels of CDKN1A by RT-qPCR in the indicated A549ACE2 or A549ACE2 TP53ko pInducer20 cells treated with or without 1 μg/mL doxycycline (dox) for 48 hr (mean ± s.e.m.). The Welch two-sample t-test with Benjamini-Hochberg p-value correction was used to assess statistical significance relative to the EGFP -Dox condition: ***, p < 0.001; **, p < 0.01; *, p < 0.05; n.s., not significant. F. Senescence-associated beta-galactosidase activity in the indicated A549ACE2 or A549ACE2 TP53ko pInducer20 cells treated with or without 1 μg/mL Dox for 48 hr (n = 14, mean ± s.e.m.). The Welch two-sample t-test with Holm p-value correction was used to assess statistical significance: **, p < 0.01; n.s., not significant. G. Left, ATAC-seq volcano plots of ancestral or delta spike versus EGFP expression in A549ACE2 pInducer20 cells 48 hr after 1 μg/mL doxycycline induction. Right, number of significant (FDR < 0.05) ATAC-seq peaks upon doxycycline-induced expression (48 hr, 1 μg/mL) of SARS-CoV-2 ancestral or delta spike relative to expression of EGFP in either A549ACE2 or A549ACE2 TP53^KO^ pInducer20 cells. Black triangles, peaks with increasing accessibility. Red circles, peaks with decreasing accessibility. H. Left, heatmap of relative levels of differentially expressed microRNAs (FDR < 0.05 and baseMean > 10) from ancestral or delta spike versus EGFP expression in A549ACE2 or A549ACE2 TP53ko pInducer20 cells 48 hr after 1 μg/mL doxycycline induction. Right, number of significant (FDR < 0.05) differentially expressed microRNAs upon doxycycline-induced expression (48 hr, 1 μg/mL) of ancestral or delta spike versus EGFP expression in A549ACE2 or A549ACE2 TP53ko pInducer20 cells. Black triangles, significantly upregulated microRNAs. Red circles, significantly downregulated microRNAs. I. Heatmap of relative levels of cytokines in supernatants of A549ACE2 or A549ACE2 TP53ko pInducer20 cells treated with 1 μg/mL doxycycline for 48 hr. The Welch two-sample t-test with Benjamini-Hochberg p-value correction was used to assess statistical significance. Sig, FDR < 0.05. J. Representative brightfield images of HNEpC-ACE2 pInducer20 cells transduced with the indicated gene constructs and treated with or without 1 μg/mL Dox for 48 hr. Scale bar, 200 μm. K. Western blotting analysis of supernatants of HNEpC-ACE2 pInducer20 cells transduced with the indicated gene constructs and treated with or without 1 μg/mL Dox for 48 hr.

We performed RNA-seq analysis of both ancestral and delta spike-expressing A549ACE2 cells with or without TP53 knockout. As expected, we observed significant positive enrichment of TP53 target gene signatures upon spike expression that is dependent on TP53 status (**Fig. 5C**). We also observed pathways associated with lipopolysaccharide and TNF treatment response, both of which stimulate inflammation, with increased pathway associations upon delta spike expression versus ancestral spike. Strikingly, the delta spike and not ancestral spike enriches for cell cycle, apoptosis, and senescence pathways in a TP53-dependent manner, indicative of key differences between spike variants (**Fig. 5C**). We observed similar enrichment of these pathways from VeroE6 transcriptomic profiles upon delta and alpha spike and not ancestral spike expression (**Fig. S6A**).

In light of our transcriptomic findings, we aimed to explore these spike-mediated cellular phenotypes further. We found that both delta and, to a lesser extent, ancestral spike expression in A549ACE2 cells decrease cell viability, effects which are attenuated in the absence of TP53 (**Fig. 5D**). In line with this observation, we observe an increase in CDKN1A/p21, a transcriptional target of TP53 that regulates cell cycle progression, in both A549ACE2 and VeroE6 cells upon spike expression (**Fig. 5B, 5E, S4G**)^86,87^. Moreover, we found senescence-associated beta-galactosidase activity to be increased in delta spike-expressing A549ACE2 cells but not ancestral spike-expressing cells in a TP53-dependent manner (**Fig. 5F**)^88^. Critically, the delta spike did not decrease cell viability nor increase senescence-associated beta-galactosidase activity in A549 cells without exogenous ACE2 expression (**Fig. S6B-C**).

As the activation of senescence may be associated with changes to chromatin state and microRNA expression^88^, we tested whether differences in the SARS-CoV-2 spike modulate these changes in a TP53-dependent manner. Indeed, the delta spike leads to widespread changes in chromatin accessibility as compared with the ancestral spike, changes which are largely lost in the absence of TP53 (**Fig. 5G**). This finding is concordant with the increases in chromatin accessibility we observed in VeroE6 cells expressing increasingly fusogenic spike variants (**Fig. S4C**). We also observed differential expression of multiple microRNAs associated with TP53 activation and its regulation such as miR-34b and miR-34c (**Fig. 5H**)^88,89^. Loss of TP53 dramatically reduces the breadth of spike-induced microRNA expression, further supporting the role of TP53 in the activation of host senescence.

As cellular senescence may promote the release of inflammatory cytokines via activation of the senescence-associated secretory phenotype (SASP)^90^, we assessed whether spike expression alone induces extracellular cytokine release in A549ACE2 cell culture supernatants. By cytokine array analysis, we detected CCL2/MCP1, CXCL1/GROα, and CXCL8/IL8 in cell culture supernatants upon delta spike expression in a TP53-dependent manner; however, ancestral spike expression does not promote substantial release of these cytokines (**Fig. 5I**). We were able to validate increases in CXCL1 and CXCL8 upon delta spike expression by Western blotting analysis (**Fig. 5B**). In delta-infected COVID-19 patient lung biopsies, we observed CXCL8 positive staining in proximity with TP53 signal by immunohistochemistry (**Fig. 3K**). Finally, we sought to assess whether differences in spike-induced cell-cell fusion promote differential inflammatory cytokine release in a normal airway cell context. We engineered human nasal epithelial cells expressing ACE2 (HNEpC-ACE2) and containing doxycycline-inducible ancestral and delta spike sequences, both of which induce cell-cell fusion in these cells (**Fig. 5J**). Expression of the delta spike but not ancestral spike induces the release of not only CXCL8/IL8 but also IL6, a SASP cytokine and a strong predictor of COVID-19 severity (**Fig. 5K**)^91^. Altogether, our findings suggest that spike-induced TP53 stabilization promotes the activation of senescence and the SASP, which varies with the propensity of the spike sequence to form syncytia.

### Differences in SARS-CoV-2 spike sequences modify TP53 stabilization and cell-cell fusion

Throughout the COVID-19 pandemic, numerous SARS-CoV-2 variants have emerged with varying immune evasiveness and fusogenicity. As of April 2023, the SARS-CoV-2 omicron family of viral variants continues to be the predominant strain in the United States and the United Kingdom, as well as many other areas in the world (**Fig. 6A**). Thus, we assessed how intrinsic spike fusogenicity across a panel of these variant spike sequences may modulate TP53 stabilization and observed a significant and positive correlation between cell-cell fusion and TP53 stability (**Fig. 6B-E**). As both the gamma/P.1 and omicron/BA.1 spike sequences are detectably fusogenic despite differences in protein stability (**Fig. 6C**), we tested whether infection with corresponding live viral isolates would also promote their TP53 stability. Infection at higher multiplicity of infection (MOI 0.5) with both variants leads to increased TP53 stabilization relative to mock infection (**Fig. 3I**). We note that as the omicron/BA.1 infection increases TP53 levels above gamma/P.1 levels despite similar MOI, other viral components may further modulate TP53 stabilization by the spike protein. Altogether, our findings suggest that the intrinsic fusogenicity of different SARS-CoV-2 variant spikes correlates with TP53 protein stabilization.

**Figure 6.**
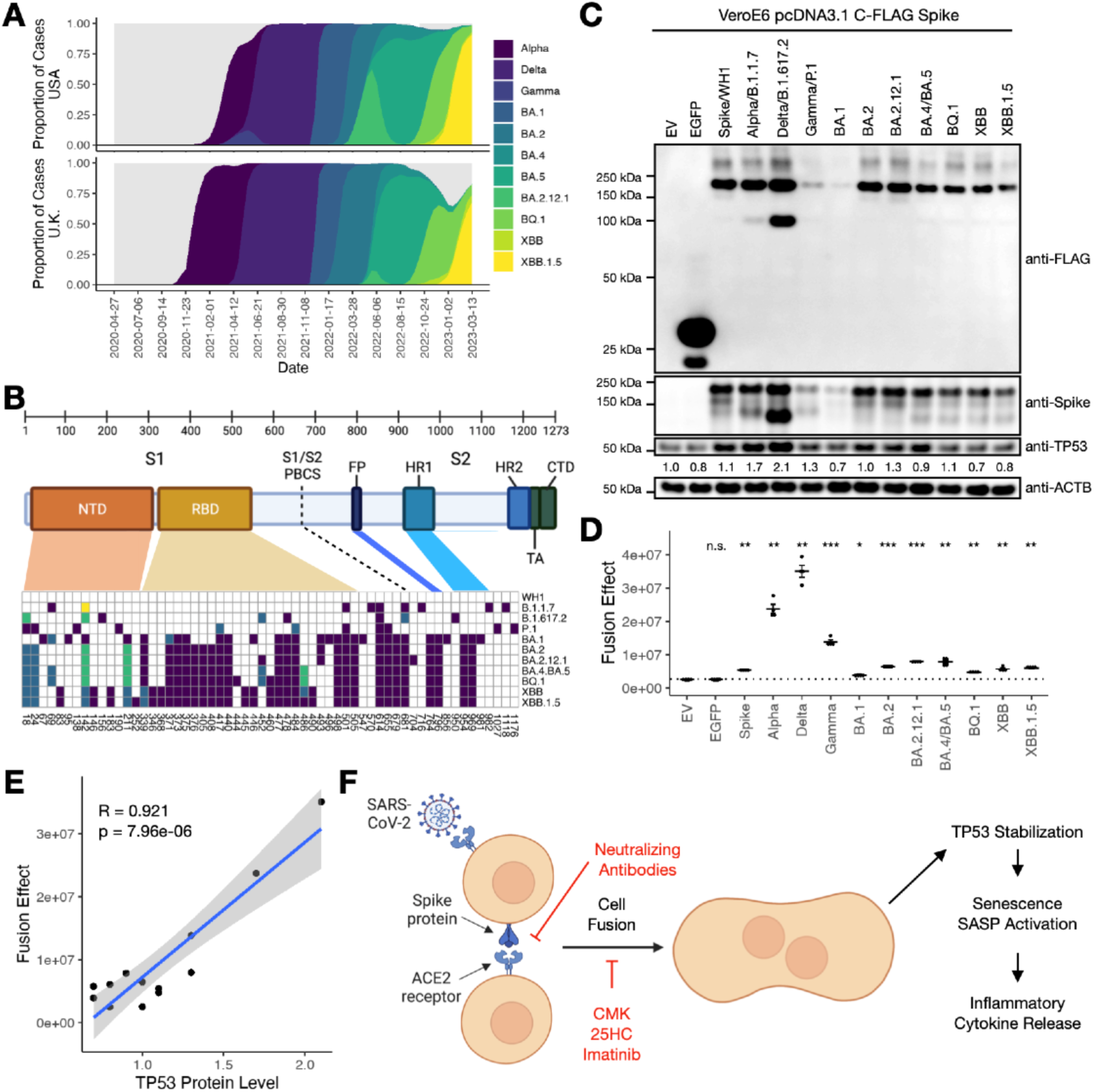
Differences in SARS-CoV-2 spike sequences modify TP53 stabilization and cell-cell fusion. A. Evolution of SARS-CoV-2 variants in the U.S.A. and the U.K. over the course of the COVID-19 pandemic. Gray, other SARS-CoV-2 variants. Data were last accessed from covariants.org on 4/3/2023^131^. B. Top, annotated SARS-CoV-2 spike subdomains. Bottom, sequence analysis of SARS-CoV-2 variant spike sequences. Colored boxes, differing sequences from ancestral spike at the given amino acid position. NTD, N-terminal domain; RBD, receptor-binding domain; PBCS, polybasic cleavage site; FP, fusion peptide; HR1/HR2, heptapeptide repeat sequences 1 and 2; TA, transmembrane anchor; CTD, C-terminal domain. C. Western blotting analysis of VeroE6 cells 48 hr after transfection with plasmids encoding the indicated FLAG-tagged proteins. EV, empty vector. Relative ratios of TP53/ACTB (normalized to EV) are annotated below TP53. D. Cell-cell fusion analysis of VeroE6 cells transfected with the corresponding spike proteins. EV, empty vector. The Welch two-sample t-test with Holm p-value correction was used to assess statistical significance relative to EV as negative control: ***, p < 0.001; **, p < 0.01; *, p < 0.05; n.s., not significant. E. Scatterplot of TP53 protein level (Fig. 6C) and fusion effect (Fig. 6D) by SARS-CoV-2 spike variant. R, Pearson correlation. p-value, Pearson correlation test. F. Proposed model of findings.

## Discussion

In summary, we performed iDAPT-MS-based chromatin proteomic analysis of host cell infection with three different pathogenic coronaviruses and identified key similarities and differences in their proteomic profiles. We unexpectedly found that TP53 protein stabilization on open chromatin occurs upon infection with SARS-CoV-2 but not MERS-CoV and HKU5-SARS-CoV-1-S and contributes to the SARS-CoV-2 host cytopathic effect. We mapped TP53 stabilization to the SARS-CoV-2 spike protein and its ability to fuse cells: increasingly fusogenic SARS-CoV-2 spike proteins increase syncytia formation and TP53 stabilization, and inhibition of syncytia formation at multiple levels attenuates TP53 levels. Critically, syncytia formation via the highly fusogenic delta spike promotes cellular senescence and extracellular cytokine release in a TP53-dependent manner. We propose a model whereby differences in syncytia formation modify the activation of TP53-dependent senescence and inflammatory cytokine release (**Fig. 6F**).

TP53 normally controls multiple host cell pathways including cell cycle, apoptosis, and senescence, and its function is modulated by viruses via a multitude of different mechanisms^50–52^. Although human coronaviruses have been previously suggested to restrict TP53 stabilization via nsp3^53,92^, the effects of SARS-CoV-2 infection on TP53 are controversial: several studies have suggested that SARS-CoV-2 decreases TP53 levels or activity^93–95^, whereas others have indicated the opposite^96–98^. We believe our findings may help explain the disparities between these studies. First, our data suggest that SARS-CoV-2 spike versus MERS-CoV and SARS-CoV-1 spikes substantially increases TP53 protein levels, which may simply overwhelm a possible nsp3-mediated TP53 inactivation mechanism. Second, several studies that have reported TP53 destabilization by SARS-CoV-2 infection used TP53-mutant Calu3 and Huh7 cell lines^94,95^. Mutant TP53 protein is often basally detected at high levels due to its inability to transactivate MDM2 expression and is degraded upon stimulation via alternate pathways, which may explain the studies’ observations^99,100^. As many mutant TP53 proteoforms retain some level of transcriptional activity, mutant TP53 may still contribute to SARS-CoV-2-mediated host cell state changes without inducing cell cycle arrest^62^.

While numerous mechanisms by which SARS-CoV-2 may induce COVID-19 inflammation have been described, one contributing mechanism is through the activation of host cellular senescence^101–105^, a stress-induced cellular state associated with irreversible cell cycle arrest, chromatin remodeling, and secretion of inflammatory cytokines via the SASP^88,106^. Indeed, CXCL8/IL8 and IL6, cytokines that correlate with severe COVID-19^91^, are associated with activation of the SASP and may precipitate the inflammatory features of severe COVID-19 even after diminution of viral levels^105^. Several mechanisms by which SARS-CoV-2 activates host cellular senescence have been reported^101,103,104,107,10895, 101^, including via SARS-CoV-2 infection directly in a TP53-dependent manner^103^. Our results implicate SARS-CoV-2 spike-induced syncytia formation as a contributing mechanism towards the TP53-dependent activation of senescence. Critically, as differences in the fusogenicity of SARS-CoV-2 spike sequences have been associated with differences in COVID-19 pathogenicity^72,109^, our model may help explain the differences in disease severity between the highly fusogenic delta/B.1.617.2 and alpha/B.1.1.7 variants relative to ancestral or omicron family viral variants^72,73,109–116^.

How does cell-cell fusion lead to TP53 stabilization? Although SARS-CoV-2 infection has been associated with activation of the DNA damage response^95,117–119^, we found that treatment with KU-55933 or berzosertib, inhibitors of the effector kinases ATM or ATR, respectively, does not substantially impact spike-induced TP53 protein stabilization (**Fig. S6D-E**)^120^. On the other hand, we found that MAPK pathway signatures are transcriptionally enriched in both A549ACE2 and VeroE6 cells expressing fusogenic spike proteins (**Fig. S6F**). MAPK pathway kinases are able to stabilize TP53 protein through its phosphorylation^121,122^; indeed, inhibition of the ERK1/2 kinases attenuates delta spike-induced TP53 stabilization in A549ACE2 cells without substantial changes in syncytia formation, suggestive of their activating roles in TP53 regulation downstream of cell-cell fusion (**Fig. S6G**). While MAPK pathway activation is a consequence of SARS-CoV-2 infection^123,124^ and can contribute to the activation of cellular senescence independent of DNA damage response^125,126^, it remains to be determined how the process of cell fusion activates the MAPK pathway.

Because the formation of syncytia by the SARS-CoV-2 spike may provoke an inflammatory response via activation of the SASP, preventing syncytia formation can be a valuable therapeutic intervention^23,24^. One approach is to use spike-targeting therapies, such as vaccines or neutralizing antibodies^127,128^. Importantly, many vaccines implement the “2P” mutant of the spike sequence, which is incapable of pathogenic membrane fusion^129^. Host-directed therapies, such as 25-hydroxycholesterol, imatinib, and CMK, may also disrupt SASP activation by inhibiting cell-cell fusion^74,81–85^. Furthermore, since ACE2 is required not only for SARS-CoV-2 viral entry but also for cell-cell fusion^20,23,78^, suppressing endogenous ACE2 expression such as through SMARCA4/2 or FXR inhibition could also prevent TP53 stabilization^48,130^.

Altogether, we propose a mechanism for how SARS-CoV-2 and its fusogenic variants differentially modulate the host cell response to infection and activate senescence via differences in the spike protein. Although the real-world immune landscape continues to change throughout the COVID-19 pandemic and cannot easily be deconvoluted from disease progression^18^, evaluation of the fusogenicity of emergent spike variants together with their effects on TP53 protein stability may facilitate the prediction of the intrinsic pathogenicity of future SARS-CoV-2 variants and nascent coronaviruses.

## Acknowledgements

We thank all members of the Slack laboratory for their input. We thank Dr. Isaac H. Solomon for providing tissue sections and clinical annotations of COVID-19 patient samples, Dr. Sidney Moyer and Dr. William Hahn for A549 TP53ko cells, Dr. Vincent Munster for SARS-CoV-1, SARS-CoV-2, and MERS-CoV spike plasmids, Dr. Benhur Lee for VSVpp virus, and Anan Quan and Dr. Adrienne Vancura for their input and support. We are grateful to the Harvard Medical School Biopolymers Facility, Harvard Medical School Research Computing, and BIDMC Confocal Imaging Core for their assistance and support. We also thank Dr. Steven P. Gygi and the Taplin Mass Spectrometry Facility at Harvard Medical School for use of their mass spectrometers and access to their COMET based data analysis suite.

## Author Contributions

J.D.L., C.B.W., F.J.S. conceived of and planned experiments. J.D.L., B.M., M.M., M.M.U., S.M.H, J.W., T.S., M.M.A., R.B.F., N.K., J.A.P. performed experiments. J.D.L., A.M., R.R.P., P.C., Y.J.H., J.A.P. performed bioinformatic analyses. J.D.L., T.M., J.A.P., C.B.W., F.J.S. provided supervision. J.D.L. and F.J.S. wrote the manuscript; all other authors reviewed and revised the manuscript.

## Supplementary Figure Legends

**Supplementary Fig 1.**
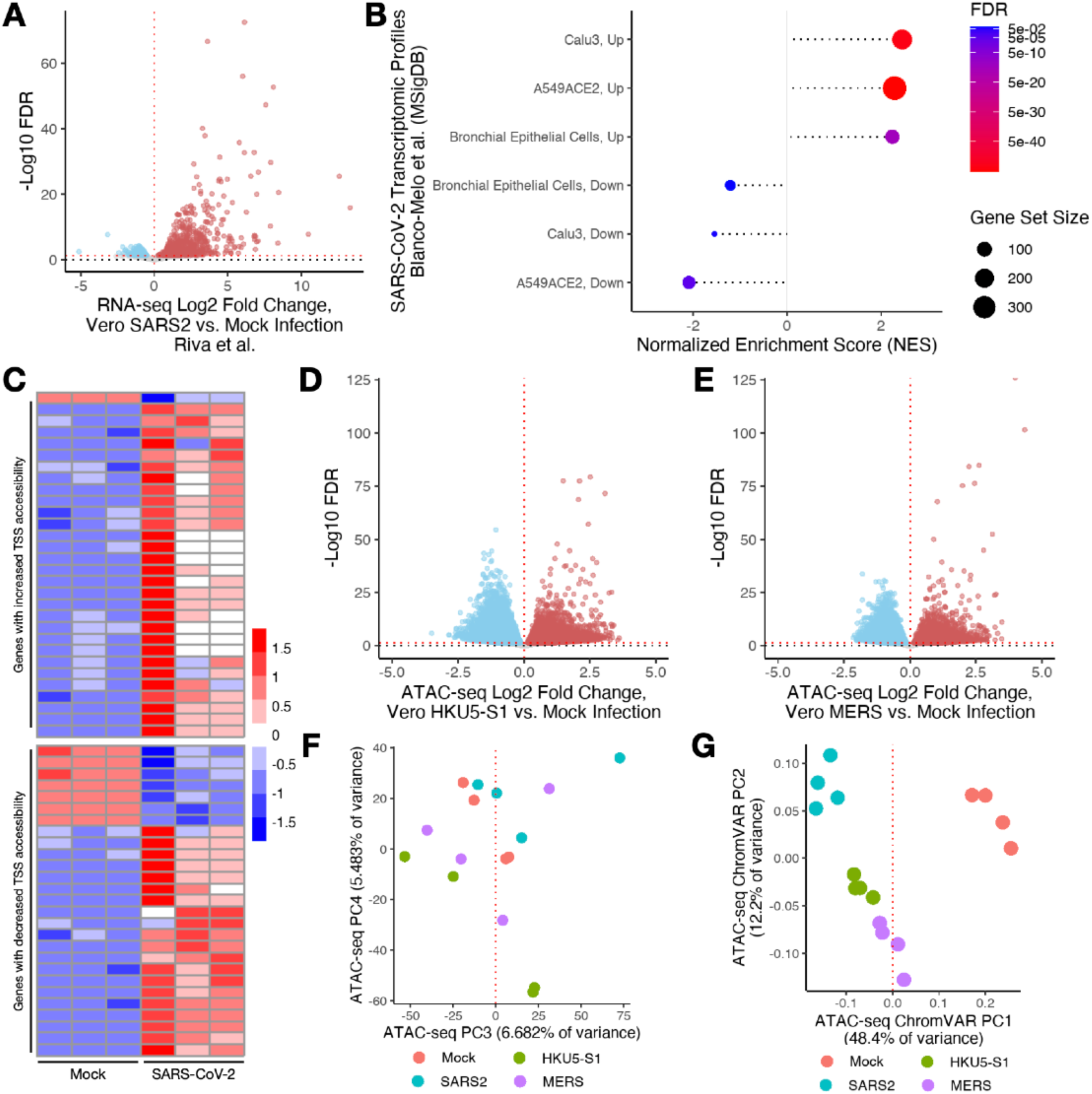
Different pathogenic coronaviruses promote different changes to host cell states. A. RNA-seq volcano plot of SARS-CoV-2 infection versus mock infection in VeroE6 cells (Riva et al.^33^). FDR, false discovery rate. B. Gene set enrichment analysis of VeroE6 transcriptome changes upon SARS-CoV-2 infection and other cell models of SARS-CoV-2 infection (Blanco-Melo et al.^34^). C. Heatmap of relative expression levels (scaled counts via the variance stabilized transform [VST]) of differentially expressed (FDR < 0.05) genes upon VeroE6 SARS-CoV-2 infection by RNA-seq and differentially accessible transcription start sites (TSS’s) (FDR < 0.05 and |log2 Fold Change| > 0.5) by ATAC-seq. Legend, Z-scaled VST counts. D. ATAC-seq volcano plot demonstrating the changes in accessible chromatin loci upon HKU5-SARS-CoV-1-S infection versus mock infection in VeroE6 cells 48 hr post-infection. FDR, false discovery rate. E. ATAC-seq volcano plot demonstrating the changes in accessible chromatin loci upon MERS-CoV infection versus mock infection in VeroE6 cells 48 hr post-infection. FDR, false discovery rate. F. Principal component analysis of ATAC-seq profiles (principal components 3 and 4). G. Principal component analysis of ChromVAR transcription factor motif profiles from ATAC-seq datasets.

**Supplementary Fig 2.**
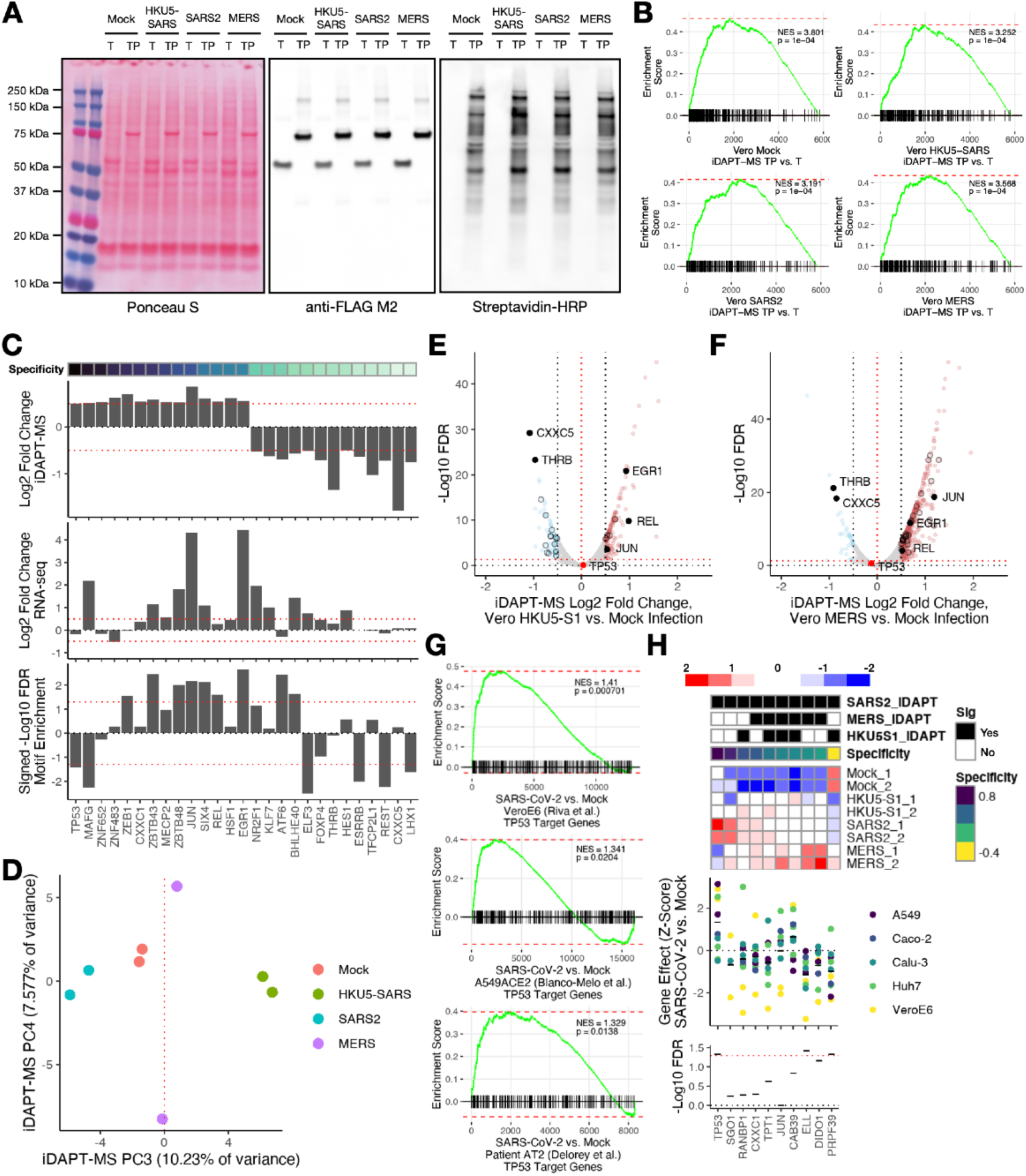
Different pathogenic coronaviruses induce different chromatin proteomic compositions upon host cell coronavirus infection. A. Representative Ponceau S and western blotting analysis of iDAPT-MS experiment. TP, transposase-peroxidase; T, transposase only. B. Gene set enrichment analysis (GSEA) of sequence-specific transcription factors of iDAPT-MS profiles comparing transposase-peroxidase (TP) enrichment versus transposase (T)- only enrichment. Log2 fold change values were used to rank proteins. NES, normalized enrichment score. p, GSEA p-value. C. Top, significant (FDR < 0.05 and |log2 Fold Change| > 0.5) transcription factor chromatin abundance changes from SARS-CoV-2 vs. mock iDAPT-MS, ordered by SARS-CoV-2 specificity. Middle, corresponding RNA abundance changes from RNA-seq. Bottom, corresponding ChromVAR motif enrichment changes from ATAC-seq. FDR, false discovery rate. D. Principal component analysis of iDAPT-MS profiles (principal components 3 and 4). E. iDAPT-MS volcano plot of HKU5-SARS-CoV-1-S infection (MOI 0.5) versus mock infection in VeroE6 cells 48 hr post-infection. Black circles, significant transcription factors (FDR < 0.05 and |log2 Fold Change| > 0.5). Black points, labeled transcription factors. Red point, TP53. F. iDAPT-MS volcano plot of MERS-CoV infection (MOI 0.5) versus mock infection in VeroE6 cells 48 hr post-infection. Black circles, significant transcription factors (FDR < 0.05 and |log2 Fold Change| > 0.5). Black points, labeled transcription factors. Red point, TP53. G. Gene set enrichment analysis of TP53 target genes (Fischer et al.^87^) in published RNA-seq (Riva et al.^33^ or Blanco-Melo et al.^34^) or single-cell RNA-seq (Delorey et al.^39^) datasets. Genes are ranked by log2 fold change. NES, normalized enrichment score. H. Top, heatmap of relative protein abundance levels of differentially abundant (FDR < 0.05 and |log2 Fold Change| > 0.5) proteins from iDAPT-MS analyses that contribute to the SARS-CoV-2 cytopathic effect (CRISPR/Cas9 pooled screening gene effect |Z| > 2) and not the MERS-CoV nor HKU5-SARS-CoV-1-S cytopathic effects (|Z| < 2). Specificity, Pearson correlation between iDAPT-MS abundances and SARS-CoV-2 infection (see Methods). Sig, significance (FDR < 0.05 and |log2 Fold Change| > 0.5) of infection condition relative to mock-infected cells. Middle, sgRNA enrichment across published CRISPR/Cas9 genetic screens (Rebendenne et al^54^). The mean Z-score of each gene is demarcated by a horizontal line. Bottom, significance of Z-score distribution per gene. The Welch one-sample t-test with Benjamini-Hochberg p-value correction was used to assess statistical significance. FDR, false discovery rate.

**Supplementary Fig 3.**
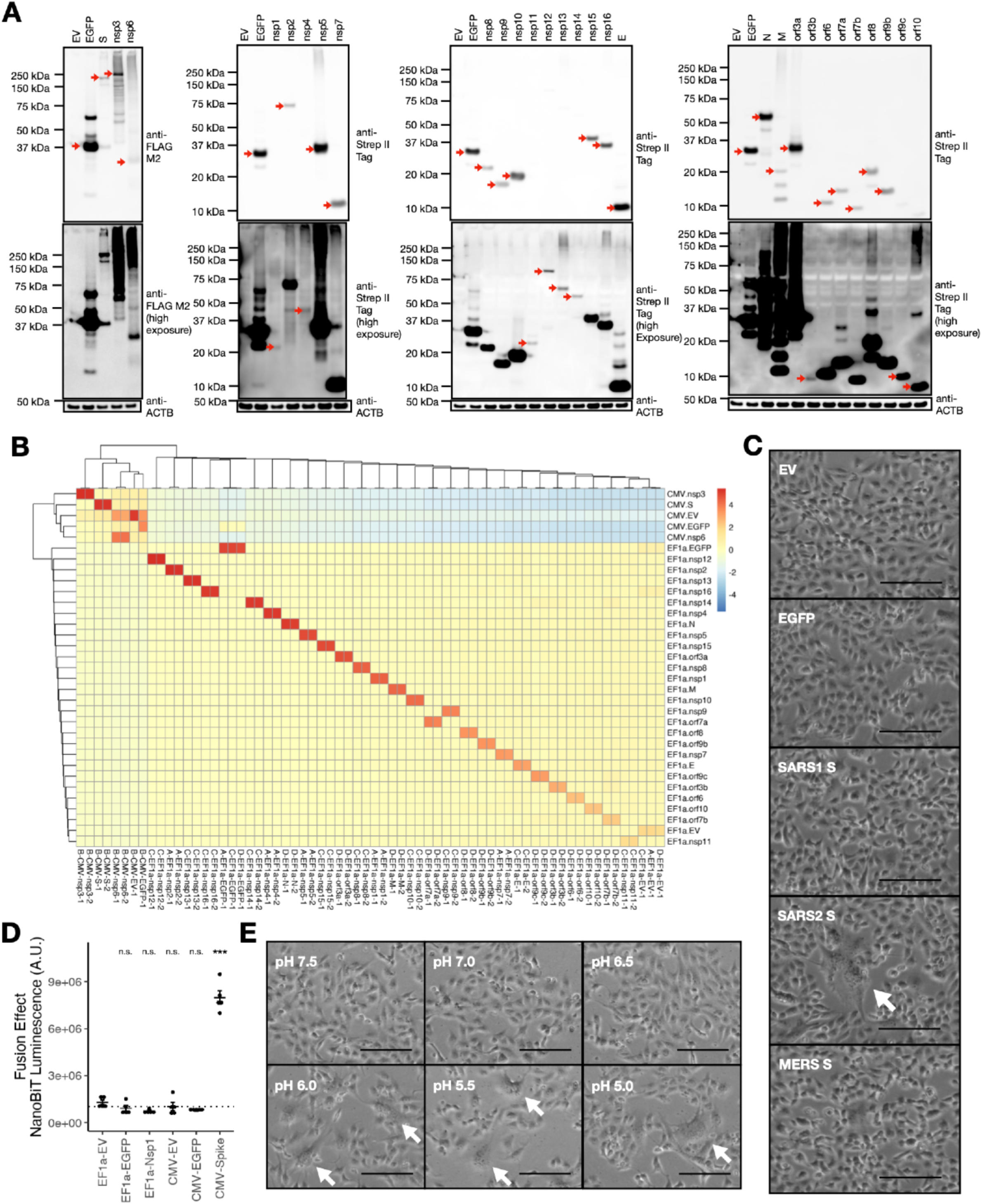
Intracellular expression of SARS-CoV-2 viral proteins leads to global chromatin accessibility changes in VeroE6 cells. A. Western blotting analysis of the indicated tagged proteins 48 hr after plasmid transfection in VeroE6 cells. B. Heatmap of sequence enrichment of aligned ATAC-seq reads to plasmid sequences. Legend, number of aligned reads Z-scaled across reference plasmid sequences. C. Representative brightfield images of VeroE6 cells transfected with plasmids encoding for the denoted genes for 48 hr. EV, empty vector. Scale bar, 200 μm. D. Cell-cell fusion analysis of VeroE6 cells transfected with the corresponding spike proteins. EV, empty vector. The Welch two-sample t-test with Holm p-value correction was used to assess statistical significance relative to EF1a-EV as negative control: ***, p < 0.001; n.s., not significant. E. Representative brightfield images of VeroE6 cells transfected with VSV-G and acidified as described in Fig. 2I. Scale bar, 200 μm.

**Supplementary Fig 4.**
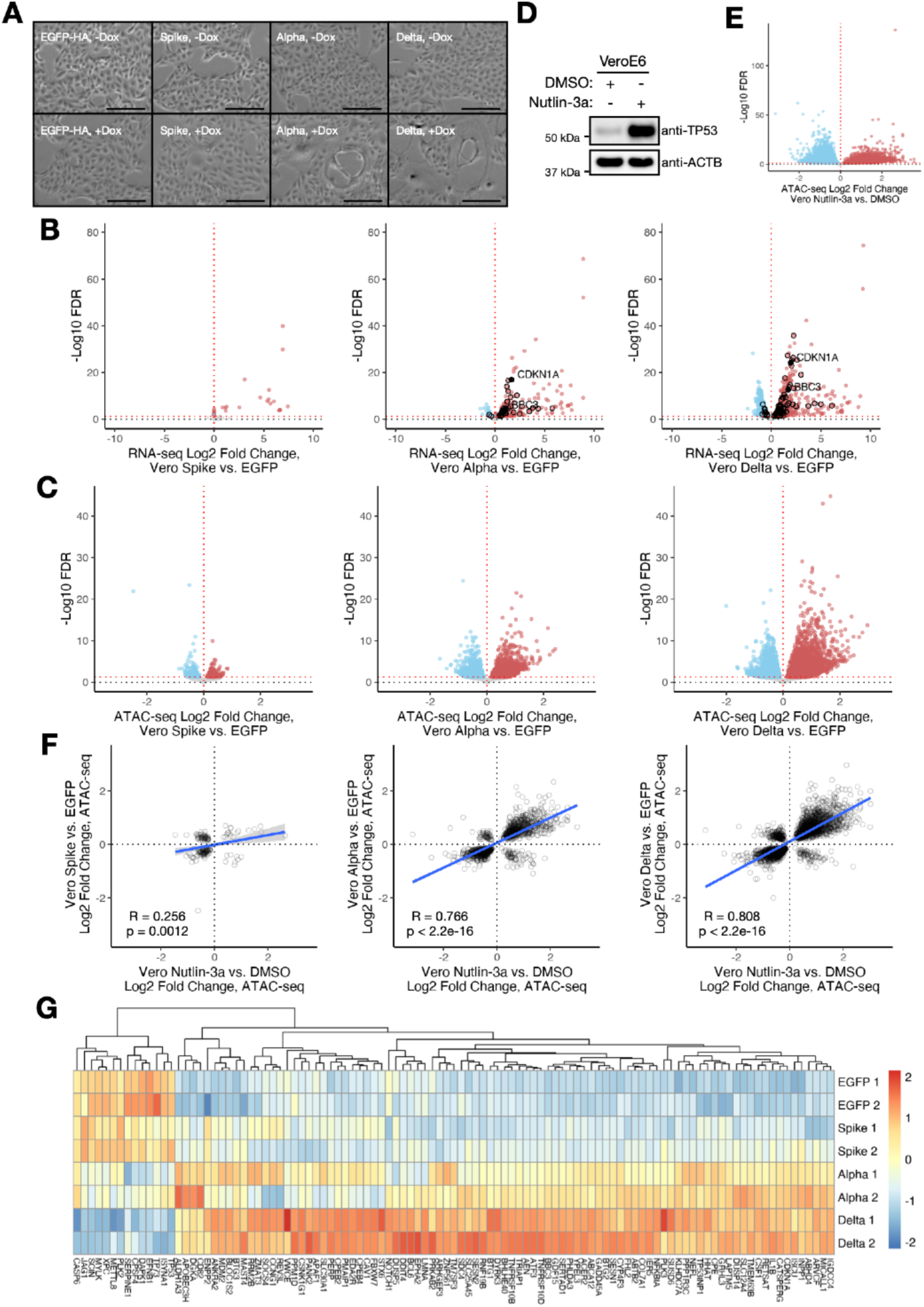
Multilevel chromatin analysis nominates TP53 as a key determinant of the host response to the expression of differentially fusogenic spike sequences. A. Representative brightfield images of VeroE6 pInducer20 cells transduced with the indicated gene constructs and treated with or without 1 μg/mL doxycycline for 48 hr. Scale bar, 200 μm. B. RNA-seq volcano plots of ancestral, alpha, or delta spike expression versus EGFP expression in VeroE6 pInducer20 cells 48 hr after 1 μg/mL doxycycline induction. Black points, significant TP53 target genes. C. ATAC-seq volcano plots of ancestral, alpha, or delta spike expression versus EGFP expression in VeroE6 pInducer20 cells 48 hr after 1 μg/mL doxycycline induction. D. Western blotting analysis of VeroE6 cells treated with either DMSO or 1 μM nutlin-3a for 24 hr. E. ATAC-seq volcano plot of 1 μM nutlin-3a versus DMSO treatment in VeroE6 cells for 24 hr. FDR, false discovery rate. F. Comparison of thresholded (FDR < 0.05) differential ATAC-seq peaks between nutlin-3a treatment (24 hr, 1 μM) and doxycycline-induced expression (48 hr, 1 μg/mL) of SARS-CoV-2 ancestral, alpha, or delta spike relative to expression of EGFP in VeroE6 pInducer20 cells. Blue line, best-fit trendline. R, Pearson correlation. p-value, Pearson correlation test. G. Heatmap of relative expression levels (scaled counts via the variance stabilized transform [VST]) of differentially expressed (FDR < 0.05) TP53 target genes from VeroE6 pInducer20 EGFP, ancestral (Spike), Alpha (B.1.1.7), or Delta (B.1.617.2) spike sequences after 48 hr of 1 μg/mL doxycycline induction. Legend, Z-scaled VST counts.

**Supplementary Fig 5.**
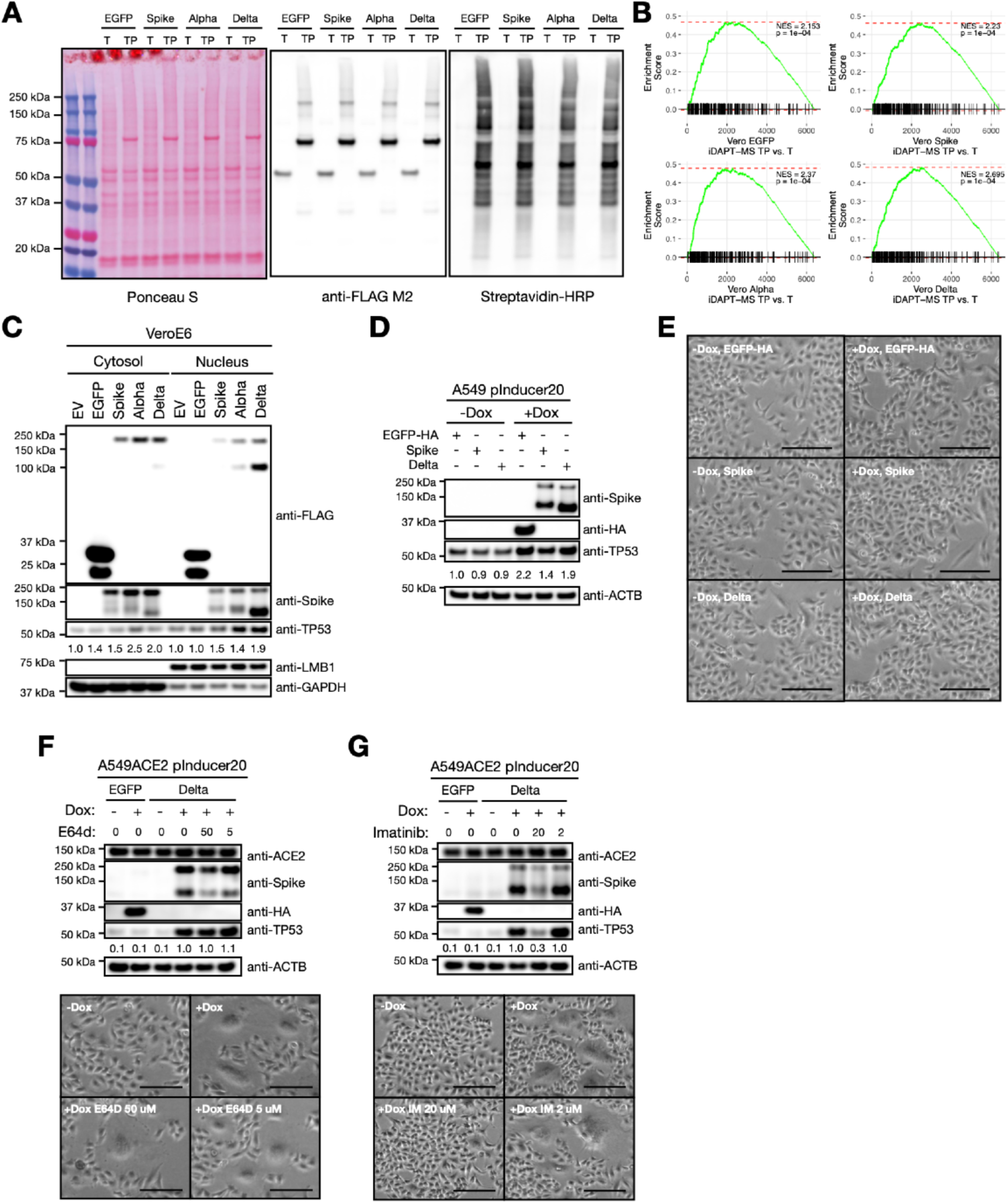
TP53 activation is a consequence of spike-induced syncytia formation. A. Representative Ponceau S and western blotting analysis of iDAPT-MS experiment. TP, transposase-peroxidase; T, transposase only. B. Gene set enrichment analysis (GSEA) of sequence-specific transcription factors of iDAPT-MS profiles comparing transposase-peroxidase (TP) enrichment versus transposase (T)- only enrichment. Log2 fold change values were used to rank proteins. NES, normalized enrichment score. p, GSEA p-value. C. Western blotting analysis of cytosolic and nuclear fractions of VeroE6 cells transfected with the indicated C-terminally FLAG-tagged proteins 48 hr after plasmid transfection. Relative ratios of TP53/GAPDH (cytosolic) or TP53/LMB1 (nuclear) (normalized to EV) are annotated below TP53. D. Western blotting analysis of A549 pInducer20 cells transduced with the indicated gene constructs and treated with or without 1 μg/mL Dox for 48 hr. Relative ratios of TP53/ACTB (normalized to EGFP -Dox) are annotated below TP53. E. Representative brightfield images of A549 pInducer20 cells transduced with the indicated gene constructs and treated with or without 1 μg/mL doxycycline for 48 hr. Scale bar, 200 μm. F. Top, Western blotting analysis of A549ACE2 pInducer20 cells transduced with the indicated gene constructs and treated with or without 1 μg/mL doxycycline (Dox) and with the corresponding amount of E64d (μM) for 24 hr. Relative ratios of TP53/ACTB (normalized to Delta +Dox) are annotated below TP53. Bottom, representative brightfield images of A549ACE2 pInducer20-Delta spike cells treated with or without 1 μg/mL doxycycline and with the corresponding amount of E64d (μM) for 24 hr. Scale bar, 200 μm. G. Top, Western blotting analysis of A549ACE2 pInducer20 cells transduced with the indicated gene constructs and treated with or without 1 μg/mL doxycycline (Dox) and with the corresponding amount of imatinib (μM) for 24 hr. Relative ratios of TP53/ACTB (normalized to Delta +Dox) are annotated below TP53. Bottom, representative brightfield images of A549ACE2 pInducer20-Delta spike cells treated with or without 1 μg/mL doxycycline and with the corresponding amount of imatinib (IM, μM) for 24 hr. Scale bar, 200 μm.

**Supplementary Fig 6.**
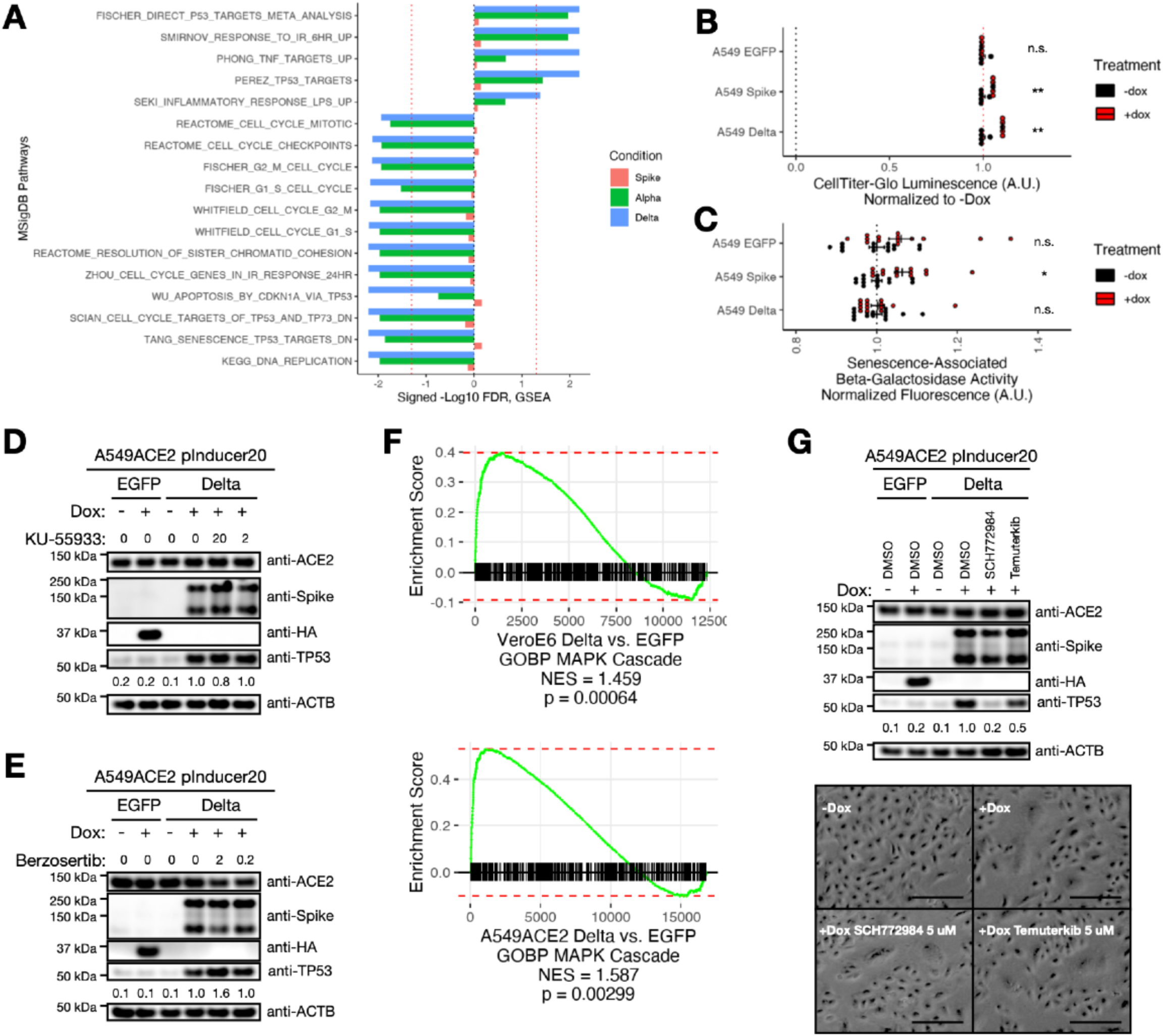
Different fusogenic SARS-CoV-2 spike sequences promote different changes to host cell states. A. MSigDB pathway enrichment analysis of differential RNA-seq profiles upon doxycycline-induced expression (48 hr, 1 μg/mL) of SARS-CoV-2 ancestral, alpha, or delta spike relative to expression of EGFP in VeroE6 pInducer20 cells. B. Cell viability as assessed by CellTiter-Glo of the indicated A549 pInducer20 cells treated with or without 1 μg/mL doxycycline (dox) for 72 hr (mean ± s.e.m.). The Welch two-sample t-test with Holm p-value correction was used to assess statistical significance: **, p < 0.01; n.s., not significant. C. Senescence-associated beta-galactosidase activity in the indicated A549 pInducer20 cells treated with or without 1 μg/mL doxycycline (dox) for 48 hr (mean ± s.e.m.). The Welch two-sample t-test with Holm p-value correction was used to assess statistical significance: *, p < 0.05; n.s., not significant. D. Western blotting analysis of A549ACE2 pInducer20 cells transduced with the indicated gene constructs and treated with or without 1 μg/mL doxycycline (Dox) and with the corresponding amount of KU-55933 (μM) for 24 hr. Relative ratios of TP53/ACTB (normalized to Delta +Dox) are annotated below TP53. E. Western blotting analysis of A549ACE2 pInducer20 cells transduced with the indicated gene constructs and treated with or without 1 μg/mL doxycycline (Dox) and with the corresponding amount of berzosertib (μM) for 24 hr. Relative ratios of TP53/ACTB (normalized to Delta +Dox) are annotated below TP53. F. Enrichment of GOBP MAPK Cascade genes in RNA-seq profiles upon expression of SARS-CoV-2 delta spike versus EGFP in either VeroE6 or A549ACE2 pInducer20 cells. Genes are ranked by log2 fold change. NES, normalized enrichment score. p-value, gene set enrichment analysis. G. Top, Western blotting analysis of A549ACE2 pInducer20 cells transduced with the indicated gene constructs and treated with or without 1 μg/mL doxycycline (Dox) and with 5 μM of SCH772984 or temuterkib for 24 hr. Relative ratios of TP53/ACTB (normalized to Delta +Dox) are annotated below TP53. Bottom, representative brightfield images of A549ACE2 pInducer20-Delta spike cells treated with or without 1 μg/mL doxycycline and with 5 μM of SCH772984 or temuterkib for 24 hr. Scale bar, 200 μm.

## Materials and Methods

### Cell lines and culture conditions

HEK293T (ATCC, CRL-3216), VeroE6 (ATCC, CRL-1586), and Huh7.5 (Washington University in St. Louis) cells were maintained in DMEM (Gibco) supplemented with 10% FBS (Sigma) and 1% penicillin/streptomycin (Thermo). A549 (ATCC, CCL-185) and A549 TP53ko cells^132^ (a gift from Dr. William Hahn) were cultured in DMEM/F-12 (Gibco) supplemented with 10% FBS (Sigma) and 1% penicillin/streptomycin (Thermo). HNEpC cells (PromoCell, C-12620) were maintained with Airway Epithelial Cell media (PromoCell, C-21160). Cells were incubated at 37 °C and 5% CO_2_.

Doxycycline (Sigma, D9891) induction of gene expression was performed with 1 μg/mL treatment in media. E64d (Selleckchem, S7393), CMK (Cayman Chem, 14965), nutlin-3a (Selleckchem, S8059), imatinib (Sigma, SML1027-10MG), berzosertib (Selleckchem, S7102), KU-55933 (Selleckchem, S1092), temuterkib (Selleckchem, S8534), and SCH772984 (MedChemExpress, HY-50846) were diluted in DMSO and added to cells at the indicated concentrations. 25-hydroxycholesterol (Sigma, H1015-10MG) was diluted in ethanol and added to cells at the indicated concentrations. SARS-CoV-2 (2019-nCoV) Spike Neutralizing Antibody (Sino Biological, 40592-R001) was diluted in water and added to cells at the indicated concentrations.

### Lentivirus production and stable cell line transduction

Lentivirus was produced in HEK293T cells via transfection of psPAX2 (Addgene #12260), VSV-G (Addgene #8454), and lentiviral transfer plasmids using standard protocols. VeroE6 cells were transfected with lentiCRISPRv2 plasmid encoding either a noncoding sgRNA (AAGGGCGCAACACTACCATT) or TP53-targeting sgRNAs (TP53-1: GGTACAGTCAGAGCCAACCT; TP53-2: TCCTCAACATCTTATCCGAG). Transfected cells were treated with 5 μg/mL puromycin (Thermo) for three days. TP53 knockout was further enriched by treatment with 10 μM nutlin-3a and validated by Western blotting analysis.

A549, A549 TP53ko, and HNEpC cells were transduced with pLEX307-ACE2-blast (Addgene #158449) to generate A549ACE2, A549ACE2 TP53ko, and HNEpC-ACE2 cell lines. VeroE6, A549, A549ACE2, A549ACE2 TP53ko, and HNEpC-ACE2 cell lines were transduced with pInducer20 lentivirus encoding EGFP, SARS-CoV-2 spike, B.1.1.7 SARS-CoV-2 spike, or B.1.617.2 SARS-CoV-2 spike sequences. Cell lines were selected with 10 μg/mL blasticidin (Invivogen) and/or 1 mg/mL geneticin (Thermo).

### VSV pseudotyped viral particle (VSVpp) production

Plasmids encoding the SARS-CoV-1, SARS-CoV-2, or MERS-CoV spike proteins were transfected in HEK293T cells. The next day, VSVp-Rluc-VSVG virus (a gift from Dr. Benhur Lee) was added to the cells for 1 hr, and then media containing anti-VSV-G (1:10,000 concentration) was added. Viral supernatant 24 hr post-infection was concentrated, aliquoted for storage, and titrated to the same luminosity in either VeroE6 or A549ACE2 cells.

Spike-mediated viral entry was determined as follows. 1e4 VeroE6 cells were seeded in 100 μL total volume in each well of a 96-well plate. The following day, 10 μL VSVpp was added and incubated for 24 hr. Luciferase activity was measured using a Glomax plate reader using the Renilla Luciferase Assay System (Promega) following the manufacturer’s instructions. Spike-dependent entry was measured as a ratio of VSVpp luminescence relative to corresponding VSVpp-VSV-G as reference.

### Coronavirus stocks

To generate viral stocks, Huh7.5 (for SARS-CoV-2) or VeroE6 (for HKU5-SARS-CoV-1-S and MERS-CoVs) were inoculated with HKU5-SARS-CoV-1-S (BEI Resources #NR-48814), SARS-CoV-2 isolate USA-WA1/2020 (BEI Resources #NR-52281), or MERS-CoV (BEI Resources #NR-48813) at a MOI of approximately 0.01 for three days to generate a P1 stock. The P1 stock was then used to inoculate VeroE6 cells for three days at approximately 50% cytopathic effects. Supernatant was harvested and clarified by centrifugation (450 *g* x 5 min), filtered through a 0.45-micron filter, and aliquoted for storage at -80°C. Viral titer was determined by plaque assay using VeroE6 cells. All work with infectious virus was performed in a Biosafety Level 3 laboratory and approved by the Yale University Biosafety Committee.

### Plasmid cloning

Gene inserts from pDONR221-EGFP (Addgene #25899), pDONR207 SARS-CoV-2 nsp3 (Addgene # 141257), pDONR223 SARS-CoV-2 NSP6 (Addgene #141260), and pDONR223 SARS-CoV-2 spike (Addgene #149329) were subcloned into pLVpuro-CMV-N-3xFLAG (Addgene #123223) by Gateway cloning. pLVX-EF1a-IRES-puro empty vector was generated by swapping the eGFP insert of pLVX-EF1alpha-eGFP-2xStrep-IRES-Puro (Addgene #141395) with a multiple cloning site. pLVX-EF1a-SARS-CoV-2-nsp16-IRES-puro was generated by subcloning the nsp16 coding sequence from pDONR223 SARS-CoV-2 NSP16 (Addgene #141269) into pLVX-EF1a-IRES-puro empty vector. All other SARS-CoV-2 viral protein-encoding plasmids were obtained from Addgene in the pLVX-EF1a-IRES-puro backbone (Addgene #141367-141370, 141372-141381, 141383-141395).

The pcDNA3.1-puro empty vector was cloned by replacing the coding sequence of pcDNA3.1 puro Nodamura B2 (Addgene #17228) with the pcDNA3.1(+) multiple cloning site. EGFP-encoding pcDNA3.1-puro was subcloned from pLVX-EF1alpha-eGFP-2xStrep-IRES-Puro (Addgene #141395). SARS-CoV-1 (Urbani), SARS-CoV-2, and MERS-CoV spike sequences from pcDNA3.1(+) plasmids generously provided by Dr. Vincent Munster were subcloned into the pcDNA3.1-puro backbone. 229E-CoV (VG40605-CF), HKU1-CoV (VG40021-UT), NL63-CoV (VG40604-CF), OC43-CoV (VG40607-CF), B.1.1.7 SARS-CoV-2 (VG40771-CF), and B.1.617.2 SARS-CoV-2 (VG40804-UT) spike plasmids were obtained from Sino Biological and subcloned into pcDNA3.1-puro. B.1.1.529/BA.1 SARS-CoV-2 spike plasmid (plv-spike-v11) was obtained from InvivoGen and subcloned into pcDNA3.1-puro. BA.2, BA.2.12.1, BA.4/BA.5, BQ.1, XBB, and XBB.1.5 spike protein mutations were obtained from covariants.org^131^, and corresponding gene blocks were synthesized (IDT) and assembled with the pcDNA3.1-puro backbone by HiFi DNA Assembly (NEB). All amino acid sequences were codon-optimized for expression in human cells. S-2P, D-2P, ΔPRRA, and ΔRRRA mutations were generated by Q5 site-directed mutagenesis (NEB).

Gene inserts from pDONR221-EGFP (Addgene #25899), pDONR223 SARS-CoV-2 spike (Addgene #149329), B.1.1.7 SARS-CoV-2 spike (Sino Biological, VG40771-CF), and B.1.617.2 SARS-CoV-2 spike (Sino Biological, VG40804-UT) sequences were subcloned into pInducer20 (Addgene #44012) via Gateway cloning.

pLenti-HiBiT-Blast was generated by swapping out the dCas9-VP64-T2A insert of lenti-dCas9-VP64-Blast (Addgene #61425) by Q5 site-directed mutagenesis (NEB). pLenti-LgBiT-T2A-Blast was generated by subcloning the LgBiT coding sequence from the FRB-LgBiT plasmid (Promega N2016) into the lenti-dCas9-VP64-Blast plasmid.

### Plasmid transfections

To each 60 mm plate, 1e6 VeroE6 cells were added. Cells were reverse transfected with expression plasmids via TransIT-X2 (Mirus) at a 1:3 ratio (2.5 μg plasmid, 7.5 μL TransIT-X2, and 250 μL OptiMEM). Cells were harvested 48 hr after transfection. For ATAC-seq analyses, after 8 hr post-transfection, puromycin was added at a final concentration of 5 μg/mL to enrich for positively transfected cells.

### Expression and purification of recombinant proteins

Purification of recombinant proteins was performed as previously described^31^. In brief, pTXB1-Tn5-FLAG (Addgene #160081) or pTXB1-Tn5-APEX2 (Addgene #160088) plasmids were transformed into the Rosetta2 E. coli strain (EMD Millipore) and streaked out on an LB agar plate containing carbenicillin and chloramphenicol. A single bacterial colony was inoculated into 10 mL LB with antibiotics and incubated overnight; this culture was then inoculated into 500 mL LB medium. Cultures were incubated at 37 °C until the optical density at 600 nm (OD600) reached ∼0.9. Isopropyl β-D-1-thiogalactopyranoside (IPTG) was added to a final concentration of 250 μM, cultures were incubated for 2 hr at 30 °C, and bacteria were pelleted and frozen at -80 °C.

Bacterial pellets were resuspended in 40 mL HEGX lysis buffer (20 mM HEPES-KOH pH 7.2, 1 M NaCl, 1 mM EDTA, 10% glycerol, 0.2% Triton X-100, 20 μM PMSF) and sonicated with a Sonic Dismembrator 100 (Fisher Scientific) at setting 7, with 5 pulses of 30 s on/off on ice. Lysate was spun at 15,000 x g in a Beckman centrifuge (JA-10 rotor) for 30 min at 4 °C. 1 mL 10% PEI was then added to the supernatant with agitation and clarified by centrifugation (15,000 x g, 15 min, 4 °C). Supernatant was then applied to 5 mL chitin resin (NEB) prewashed with HEGX buffer and incubated for 1 hr at 4 °C with agitation. Chitin slurry was applied to an Econo-Pak column (Bio-Rad) to remove unbound protein, washed with 20 column volumes of HEGX buffer and 1 column volume of HEGX with 50 mM DTT, and then incubated with 1 column volume of HEGX with 50 mM DTT for 48 hr at 4 °C. After elution, the column was washed with 1 column volume of 2x dialysis buffer (2xDB: 100 mM HEPES-KOH pH 7.2, 0.2 M NaCl, 0.2 mM EDTA, 20% glycerol, 0.2% Triton X-100, 2 mM DTT). Eluates were combined, concentrated with a 10 kDa MWCO centrifugal filter (EMD Millipore), and subjected to buffer exchange with 2xDB using PD-10 desalting columns (GE Healthcare). Eluates were further concentrated, and precipitates were removed by centrifugation.

Soluble proteins (Tn5, TP) were normalized by their transposase activities as described below for ATAC-seq with the following amendments. Serial dilutions of purified protein in 2xDB were added to a constant 12.5 pmol MEDS-A/B (1.25 μL in water) and incubated at room temperature for 1 hr. 100,000 K562 (ATCC) cells per condition were harvested, lysed, and tagmented with fresh transposome preparations. DNA libraries were extracted with DNA Clean and Concentrator-5 (Zymo), and optimal PCR cycle number was determined via quantitative PCR^31,36^. Transposase activity was determined as the dilution leading to the lowest PCR cycle number and similar to a reference transposase preparation performed in parallel. Normalized protein stocks were aliquoted, snap frozen with liquid nitrogen and stored at -80 °C.

### Transposome adaptor preparation

All transposome adaptors were synthesized at Integrated DNA Technologies (IDT). The oligonucleotide sequences were similar as previously described^31^: Tn5MErev, 5’- [phos]CTGTCTCTTATACACATCT-3’; Tn5ME-A, 5ʹ-TCGTCGGCAGCGTCAGATGTGTATAAGAGACAG- 3’; Tn5ME-B: 5’-GTCTCGTGGGCTCGGAGATGTGTATAAGAGACAG-3’. All oligos were resuspended in water to a final concentration of 200 μM each. Equimolar amounts of Tn5MErev/Tn5ME-A, Tn5MErev/Tn5ME-B were added together in separate tubes, denatured at 95 °C for 10 min, and cooled slowly to room temperature by removing the heat block. Tn5MEDS-A/Tn5MEDS-B was combined at equimolar amounts to form 100 μM aliquots and stored at -20 °C.

### Integrated DNA and Protein Tagging (iDAPT) analysis

On the day of harvest, VeroE6 cells were washed and suspended in 1xPBS. Cells were pelleted (500 x g, 5 min, 4 °C), incubated in 500 μL nuclei extraction buffer (20 mM HEPES-KOH pH 7.9, 10 mM KCl, 0.5 mM spermidine, 0.1% Triton X-100, 20% glycerol, and 1x EDTA-free protease/phosphatase inhibitors [Thermo] in ultrapure water) per ∼5e6 cells for 10 min on ice, and 50 μL DMSO mixed into the nuclear suspension prior to freezing cells at -80 °C. We found these lysis conditions to inactivate live SARS-CoV-2 virus for downstream handling and be compatible with high-quality ATAC-seq/iDAPT-MS profiling.

2.5 μmol MEDS-A/B, corresponding recombinant transposase (TP [transposase-peroxidase] or T [Tn5 transposase]) enzyme, and 1 μmol hemin chloride per sample were incubated at room temperature for 1 hr. Nuclei were thawed on ice, pelleted (1000 x g, 10 min, 4 °C), and resuspended in tagmentation buffer (20% dimethylformamide, 10 mM MgCl_2_, 20 mM Tris-HCl pH 7.5, 33% 1xPBS, 1% BSA, 0.01% digitonin, 0.1% Tween-20, 500 μM biotin-phenol, and 1x protease/phosphatase inhibitor) without transposome. Nuclei were counted with trypan blue (Thermo) with a Countess 3 (Thermo), and 5e6 nuclei per condition was resuspended in tagmentation buffer and 1 μmol transposome equivalent in a total volume of 500 μL, followed by incubation at 37 °C for 30 min with agitation on a thermomixer (1,000 rpm). 10 μL of tagmentation mix was saved for quality assessment as described below for ATAC-seq sample preparation. The remaining nuclear suspension was then washed 2x with 1xPBS supplemented with 500 μM biotin-phenol, 0.1% BSA, 0.1% Tween-20, and 1x protease/phosphatase inhibitor (3000 x g, 5 min, 4 °C) and labeled with 1 mM hydrogen peroxide and 500 μM biotin-phenol for 1 min in 1xPBS with 1x protease/phosphatase inhibitor in a volume of 500 μL. Peroxidation reactions were quenched with 500 μL 2x quenching buffer (10 mM Trolox, 20 mM sodium ascorbate, 20 mM NaN_3_, and 1x protease inhibitor in 1xPBS). Labeled nuclei were then pelleted, washed with 1x quenching buffer, resuspended in 500 μL RIPA (Boston BioProducts) containing protease/phosphatase inhibitors, and frozen at -80 °C. Lysates were thawed on ice, sonicated via a Sonic Dismembrator 100 (Fisher Scientific, setting 3, 15 s, 4 pulses), and incubated on ice for 1 hr after the addition of 2 μL benzonase (EMD Millipore). Lysates were clarified by centrifugation (15,000 x g, 20 min, 4 °C), quantified via BCA assay (Thermo Fisher Scientific), and subjected to either Western blotting or quantitative mass spectrometry analyses as described below.

### Western blotting analysis

Whole cell lysates were generated by resuspending cells in RIPA (Boston BioProducts) supplemented with 1x EDTA-free protease/phosphatase inhibitor cocktail (Thermo). Lysates were sonicated via a Sonic Dismembrator 100 (Fisher Scientific) at setting 3 with 3-4 pulses of 15 s on/off on ice and incubated for an additional 1 hr at 4 °C with end-to-end rotation. Lysates were clarified by centrifugation (15,000 x g, 20 min, 4 °C) and their concentrations quantified via the BCA assay (Thermo Fisher Scientific). Lysates were run on NuPAGE 4-12% Bis-Tris protein gels (Thermo Fisher Scientific) and transferred to 0.45 μm nitrocellulose membranes (BioRad). Membranes were blocked with 3% milk in PBS-T and incubated overnight with primary antibody and subsequently with secondary antibody after brief washing with PBS-T. Chemiluminescence was determined by applying ECL Western Blotting detection reagent (GE Healthcare) or SuperSignal West Femto Maximum Sensitivity Substrate (Thermo) to membranes and imaging on an Amersham Imager 600 (GE Healthcare). Membranes were stripped with Restore PLUS Stripping Buffer (Thermo Fisher Scientific). Images were processed with Fiji/ImageJ v2.3.0.

Primary antibodies used were anti-FLAG M2 (Sigma-Aldrich, F1804, 1:2000), anti-DYKDDDDK (FLAG) Tag (CST, 14793S, 1:1000), anti-ACE2 (ProSci, 3217, 1:1000), anti-ACTB (Sigma, A5316, 1:5000), anti-SARS-CoV-2-Spike (Sino Biological, 40592-MM117, 1:2000), anti-MERS-CoV-Spike (Sino Biological, 100208-RP02, 1:1000), anti-Renilla Luciferase (abcam, ab185925, 1:1000), anti-VSV-G [8G5F11] (Kerafast, EB0010, 1:1000), anti-VSV-M [23H12] (Kerafast, EB0011, 1:1000), anti-HA (CST, 2367S, 1:1000), Anti-TP53 (CST, 9282S, 1:1000), anti-CDKN1A/p21 (CST, 2947S, 1:1000), anti-CXCL8/IL8 [E5F5Q] (CST, 94407, 1:1000), anti-CXCL1/2 [E5m6D] (CST, 24376, 1:1000), anti-Lamin B1 (abcam, ab16048, 1:1000), anti-GAPDH [14C10] (CST, 2118, 1:1000), and anti-Strep II Tag (Novus, NBP2-43735, 1:1000). Secondary antibodies used were Rabbit IgG, HRP-linked F(ab’)_2_ fragment (GE Healthcare NA9340, from donkey, 1:5000) and Mouse IgG, HRP-linked whole Ab (GE Healthcare NA931, from sheep, 1:5000). Streptavidin-HRP (CST 3999S, 1:1000) was also used for probing.

### Streptavidin enrichment and tandem mass tag labeling

250 μg lysate was reduced with 5 mM DTT in 500 μL RIPA and then added to 60 μL Pierce streptavidin bead slurry equilibrated 2x with RIPA buffer. Lysate/bead mix was incubated with end-to-end rotation overnight at 4 °C. Beads were washed with RIPA 2x, 1 M KCl 1x, 0.1 M Na_2_CO_3_ (1x), 2 M urea in 10 mM Tris-HCl pH 8.0 1x, and 200 mM EPPS pH 8.5 3x prior to resuspension in 100 μL 200 mM EPPS pH 8.5, with beads resuspended and incubated with end-to-end rotation for 5 min per wash. 1 μL mass spectrometry-grade LysC (Wako) was added to each tube and incubated at 37 °C for 3 hr with mixing, and an additional 1 μL mass spectrometry-grade trypsin (Thermo Fisher Scientific) was added, followed by overnight incubation at 37 °C with mixing. Beads were magnetized, and eluate was collected and subjected to downstream TMT labeling.

Peptides were processed using the SL-TMT method^133^. TMT reagents (0.8 mg) were dissolved in anhydrous acetonitrile (40 μL), of which 10 μL was added to each peptide suspension (100 μL) with 30 μL of acetonitrile to achieve a final acetonitrile concentration of approximately 30% (v/v). Following incubation at room temperature for 1 h, the reaction was quenched with hydroxylamine to a final concentration of 0.3% (v/v). The TMT-labeled samples were pooled at equal volumes across all samples. The pooled sample was vacuum centrifuged to near dryness and subjected to C18 solid-phase extraction (SPE) (Sep-Pak, Waters).

### Off-line basic pH reversed-phase (BPRP) fractionation

We fractionated the pooled TMT-labeled peptide sample using BPRP HPLC^134^. We used an Agilent 1200 pump equipped with a degasser and a photodiode array (PDA) detector (set at 220 and 280 nm wavelength) from ThermoFisher Scientific (Waltham, MA). Peptides were subjected to a 50-min linear gradient from 9% to 35% acetonitrile in 10 mM ammonium bicarbonate pH 8 at a flow rate 600 μL/min over an Agilent 300Extend C18 column (3.5 μm particles, 4.6 mm ID and 220 mm in length). The peptide mixture was fractionated into a total of 96 fractions, which were consolidated into 24 super-fractions^135^. Samples were subsequently acidified with 1% formic acid and vacuum centrifuged to near dryness. Each consolidated fraction was desalted via StageTip, dried again via vacuum centrifugation, and reconstituted in 5% acetonitrile, 5% formic acid for LC-MS/MS processing.

### LC-MS/MS proteomic analysis

Samples were analyzed on an Orbitrap Fusion mass spectrometer (Thermo Fisher Scientific, San Jose, CA) coupled to a Proxeon EASY-nLC 1200 liquid chromatography (LC) pump (Thermo Fisher Scientific). Peptides were separated on a 100 μm inner diameter microcapillary column packed with 35 cm of Accucore C18 resin (2.6 μm, 150 Å, ThermoFisher). For each analysis, approximately 2 μg of peptides were separated using a 150 min gradient of 8 to 28% acetonitrile in 0.125% formic acid at a flow rate of 450-500 nL/min. Each analysis used an MS3-based TMT method^136,137^, which has been shown to reduce ion interference compared to MS2 quantification^138^. The data were collected as described previously using an SPS-MS3 method^139^.

### Proteomic data analysis

Mass spectra were processed using a Sequest-based pipeline^140^, as described previously^141^. Database searching included all entries from the C. sabaeus and coronavirus UniProt databases, which was concatenated with one composed of all protein sequences in the reversed order. Oxidation of methionine residues (+15.995 Da) was set as a variable modification, and TMT tags on lysine residues and peptide N-termini (+229.163 Da) and carbamidomethylation of cysteine residues (+57.021 Da) were set as static modifications. Peptide-spectrum matches (PSMs) were adjusted to a 1% false discovery rate (FDR)^142,143^ using a linear discriminant analysis (LDA), as described previously^140^. For quantitation, we extracted the summed signal-to-noise (S:N) ratio for each TMT channel and omitted PSMs with poor quality, MS3 spectra with TMT reporter summed signal-to-noise of less than 100, or isolation specificity < 0.7^144^.

A two-step analysis for data quality assessment was performed as follows. First, PSMs without further normalization were log2-transformed and collapsed to proteins by arithmetic average, with priority given to uniquely mapping peptides. Differential protein enrichment (TP vs. T) for each condition was determined using the limma package in R, and iDAPT on-target enrichment was confirmed by gene set enrichment analysis (fgsea v1.16.0 in R) of C. sabaeus sequence-specific transcription factors obtained from the CisBP v2.0 database (http://cisbp.ccbr.utoronto.ca/), using gene symbols ranked by their log2 fold changes. To filter out background signal, we used a receiver operating characteristic analysis with CisBP transcription factors as positive controls and “contaminant” or endogenously biotinylated proteins (ACACA, PC, PCCA, PCCB, MCCC1, MCCC2) as negative controls. Proteins below the optimal threshold in all our datasets were excluded from downstream analysis.

Using our filtered protein list, PSMs were log2-transformed and collapsed to proteins by arithmetic mean, with priority given to uniquely mapping peptides. Clustering analyses were performed with these post-filtered normalized protein abundances. Heatmaps were generated using pheatmap (v1.0.12). Differential protein enrichment relative to the negative control condition was determined using the limma package (v3.46.0).

### ATAC-seq sample preparation

The OmniATAC sample preparation protocol was used as previously described with modifications where indicated below^145^. Normalized transposase enzyme (in 2xDB) was mixed with 12.5 pmol MEDS-A/B (1.25 μL in water) and incubated at room temperature for 1 hr. In the meantime, 100,000 cells were centrifuged at 500 x g for 5 min at 4°C. Cells were resuspended in 50 μL lysis buffer 1 (LB1: 10 mM Tris-HCl pH 7.5, 10 mM NaCl, 3 mM MgCl2, 0.01% digitonin, 0.1% Tween-20, and 0.1% NP-40) with trituration, incubated on ice for 3 min, and then further supplemented with 1 mL lysis buffer 2 (LB2: 10 mM Tris-HCl pH 7.5, 10 mM NaCl, 3 mM MgCl2, and 0.1% Tween-20). Nuclei were pelleted (500 x g, 10 min, 4 °C), resuspended with 50 μL tagmentation reaction mixture (20% dimethylformamide, 10 mM MgCl_2_, 20 mM Tris-HCl pH 7.5, 33% 1xPBS, 0.01% digitonin, 0.1% Tween-20, and transposome complex in 50 μL total volume), and incubated at 37°C for 30 min with agitation on a thermomixer (1,000 rpm). After tagmentation, DNA libraries were extracted with DNA Clean and Concentrator-5 (Zymo) and eluted with 21 μL water.

To determine optimal PCR cycle number for library amplification, quantitative PCR was performed similarly as previously reported on a StepOnePlus Real-Time PCR (Applied Biosystems) with the StepOne v2.3 software^36^. 2 μL of each ATAC-seq library was added to 2x NEBNext Master Mix (NEB) and 0.4x SYBR Green (Thermo Fisher) with 1.25 μM of each primer (Primer 1.1: 5’- AATGATACGGCGACCACCGAGATCTACACTAGATCGCTCGTCGGCAGCGTCAGATGTGTAT-3’; Primer 2.1: 5’-CAAGCAGAAGACGGCATACGAGATTCGCCTTAGTCTCGTGGGCTCGGAGATGT-3’) in a final volume of 15 μL, and quantification was assessed using the following conditions: 72 °C for 5 min; 98 °C for 30 s; and thermocycling at 98 °C for 10 s, 63 °C for 30 s and 72 °C for 1 min. Optimal PCR cycle number was determined as the qPCR cycle yielding fluorescence between 1/4 and 1/3 of the maximum fluorescence. The remaining DNA library was then amplified accordingly by PCR using previously reported barcoded primers for library multiplexing^146^, purified with DNA Clean and Concentrator-5 (Zymo), and eluted into 20 μL final volume with water. Libraries were then subject to TapeStation 2200 High Sensitivity D1000 fragment size analysis (Agilent), MiSeq Nano v2 sequencing for library normalization, and NovaSeq 6000 paired-end sequencing for deep sequencing (Illumina).

### ATAC-seq data preprocessing

Paired-end sequencing reads were trimmed with TrimGalore v0.4.5 to remove adaptor sequence CTGTCTCTTATACACATCT, which arises at the 3’ end due to sequenced DNA fragments being shorter than the sequencing length (75 bp). For plasmid-transfected ATAC-seq datasets, trimmed reads were first aligned to plasmid DNA sequences using bowtie2 v2.2.9 with options “--no-mixed --dovetail --un-conc -X 2000”. Unaligned reads or reads from other experiments were aligned to either the hg38 or ChlSab1.1 reference genomes using bowtie2 v2.2.9 with options “--no-unal -- no-discordant --no-mixed -X 2000”. Reads mapping to the mitochondrial genome were subsequently removed, and duplicate reads were removed with Picard v2.8.0. For transcription start site (TSS) enrichment, reads were downsampled to approximately 5 million paired-end fragments. TSS enrichment was performed by first shifting insert positions aligned to the reverse strand by -5 bp and the forward strand by +4 bp as previously described^36^ and then determining the distance of each insertion to the closest Ensembl v104 transcription start site with Homer v4.9. Principal component analyses were performed with counts transformed by the varianceStabilizingTransformation function from DESeq2. Figures were generated with ggplot2 v3.3.3. Peaks were determined by MACS2 v2.1.1 using options “callpeak --nomodel --shift -100 - -extsize 200 --nolambda -q 0.01 --keep-dup all”.

ATAC-seq peaks for VeroE6 analyses were obtained by generating a consensus peak set after consolidating all reads from coronavirus- and mock-infected samples or pInducer20 samples. Peaks overlapping 500 bp were clipped with bedClip (ucsc-tools v363). Read counts for VeroE6 infection, plasmid transfection, and recombinant protein treatment ATAC-seq datasets were determined by bedtools multicov using VeroE6 virus infection consensus peaks. Count matrices were processed with DESeq2 for differential transposon insertions with shrunken log2 fold changes. Read counts for VeroE6 pInducer20 and nutlin-3a treatment ATAC-seq datasets were determined by bedtools multicov using VeroE6 pInducer20 consensus peaks. Count matrices were processed with DESeq2 for differential transposon insertions with shrunken log2 fold changes.

ATAC-seq peaks for A549ACE2 and A549ACE2 TP53ko pInducer20 analyses were obtained by generating a consensus peak set after consolidating all reads from these datasets and then excluding blacklist regions (https://github.com/Boyle-Lab/Blacklist/blob/master/lists/hg38-blacklist.v2.bed.gz) with bedtools subtract (v2.27.1). Read counts for A549ACE2 pInducer20 ATAC-seq datasets were determined by bedtools multicov. Count matrices were processed with DESeq2 for differential transposon insertions with shrunken log2 fold changes.

### ATAC-seq transcription factor motif analysis

A BSgenome package was forged using the Ensembl v104 chlSab2.2bit file and the BSgenome (v1.58.0) library using the forgeBSgenomeDataPkg function in R. CisBP 2.0 motifs corresponding to C. sabaeus were downloaded from http://cisbp.ccbr.utoronto.ca/. Motif enrichment analysis was performed with ChromVAR as previously described^147,148^. ChromVAR motif deviations from the computeDeviations function were used for principal component analysis, and FDR-adjusted p-values were obtained with the differentialDeviations function with default settings.

### CRISPR/Cas9 host cell genetic dependency analysis

CRISPR/Cas9 pooled screening datasets of VeroE6 host factors in SARS-CoV-2, MERS-CoV, and HKU5-SARS-CoV-1-S infections were obtained from Wei et al.^32^ (https://www.cell.com/cms/10.1016/j.cell.2020.10.028/attachment/8c54b670-8d32-409f-9119-9e00ba32a73c/mmc2.xlsx). CRISPR/Cas9 pooled screening datasets of host factors in SARS-CoV-2 infection across multiple cell line models were obtained from Rebendenne et al.^54^ (https://www.ncbi.nlm.nih.gov/pmc/articles/PMC8168385/bin/2a8527d91253de19a3b16993.xlsx). CRISPR/Cas9 screening studies analyzed by Rebendenne et al. include VeroE6^32,54^, A549-ACE2^55,56^, Huh7^57^ and derivatives (Huh7.5^58^ and Huh7.5.1-ACE2-TMPRSS2^59^), Caco-2-ACE2^54^, and Calu-3^54,60^. Neither VeroE6 nor A549 have reported TP53 mutations, whereas Caco-2 (E204*), Calu-3 (M237I), and Huh7 (Y220C) have homozygous mutations in TP53^61^. The Z-score per gene in each cell line or infection condition was used as is. A Z-score threshold of +/- 2 was applied to prioritize gene effects on host cell states upon coronavirus infection.

### RNA-seq library construction

Cells were lysed using Trizol reagent (Thermo Fisher) and RNA was extracted by phase separation. Total RNA was quantified by nanodrop (DeNovix DS-11) and TapeStation 4200 (RNA Screen Tape; Agilent Technologies, Inc) or 2100 Bioanalyzer (Total RNA Nano; Agilent Technologies, Inc). 500 ng of total RNA was used for library preparation via the KAPA mRNA HyperPrep kit (Roche) according to the manufacturer’s recommendations. Briefly, mature mRNA was isolated from total RNA using mRNA Capture Beads. Following elution in Fragment, Prime and Elute buffer (FPE), RNA was fragmented using heat (6 min at 94°C) in the presence of Mg^2+^. First and second strand synthesis and A-tailing were performed according to manufacturer’s recommendations. 7 μM KAPA Unique Dual-Indexed (UDI, Roche) adapters were ligated to the second strand synthesis product in the presence of a ligation master mix in a reaction that was performed at 20 °C for 15 min. Following cleanup, all libraries underwent 11 cycles of amplification. Successful library production, quality control, and quantification was assessed using Tapestation (High sensitivity D1000 Screen Tape, Agilent). Libraries were pooled together and subjected to NovaSeq 6000.

### RNA-seq analysis

Raw sequencing reads were aligned to a reference transcriptome generated from the Ensembl v104 GRCh38 or ChlSab1.1 databases with salmon v1.4.0 using options “--seqBias --useVBOpt -- gcBias --posBias --numBootstraps 30 --validateMappings”. Length-scaled transcripts per million were acquired using tximport v1.18.0, and shrunken log2 fold changes and false discovery rates were determined by DESeq2 in R. Principal component analysis and heatmap analysis were performed with counts transformed by the varianceStabilizingTransformation function from DESeq2.

TP53 target gene symbols from Fisher et al. were obtained from: https://static-content.springer.com/esm/art%3A10.1038%2Fonc.2016.502/MediaObjects/41388_2017_BFonc2016502_MOESM7_ESM.xlsx, and gene set enrichment analysis was performed using corresponding gene symbols ranked by log2 fold changes. Human Entrez gene IDs (either as is for A549ACE2 datasets or corresponding to matching C. sabaeus gene symbols) were ranked by log2 fold changes were used for gene set enrichment analysis. MSigDB C2 (https://www.gsea-msigdb.org/gsea/msigdb/genesets.jsp?collection=C2) pathways were used for pathway enrichment analysis.

Raw A549ACE2 RNA-seq datasets were obtained from Blanco-Melo et al.^34^ with accession number GSE147507 and processed as described above. VeroE6 RNA-seq read counts from an RSEM alignment pipeline were obtained from GSE153940 (Riva et al.^33^) and processed via DESeq2 analysis as described above. Single cell RNA-seq from Delorey et al.^39^ were obtained from https://static-content.springer.com/esm/art%3A10.1038%2Fs41586-021-03570-8/MediaObjects/41586_2021_3570_MOESM6_ESM.xlsx. Gene symbols were ranked by “disease_log2fc” after subsetting by AT2 cell type and pseudobulk analysis.

### Brightfield imaging

At the time of imaging, cells were washed and maintained with 1xPBS. Images were acquired via an EVOS FL Cell Imaging System (Life Technologies) at 4x magnification.

### VSV-G acidification

4e5 VeroE6 cells were plated in 6-well plates and transfected with 1 μg VSV-G plasmid (Addgene #8454) via TransIT-X2 (Mirus). After 24 hr post-transfection, wells were washed twice with PBS and then subjected to DMEM (12100046) buffered with 10 mM morpholineethanesulfonic acid (MES) at varying pH levels. Cells were washed with PBS and then incubated in cell culture media. Cells were analyzed 24 h after acidification.

### Tissue staining and immunohistochemistry

Post-mortem lung specimens from individuals with severe COVID-19 were collected through an excess tissue waived consent protocol approved by the Mass General Brigham Institutional Review Board and were provided by Dr. Isaac H. Solomon. Consent for autopsy was provided by the patients’ next of kin or healthcare proxy per Massachusetts state law. All autopsies were performed at Brigham and Women’s Hospital from 9/1/2020 to 1/30/22 with pre- or peri-mortem positive testing for SARS-CoV-2 by nasopharyngeal swab qPCR. At the time of autopsy, lung tissues were fixed in formalin and embedded in paraffin. Serial tissue sections were mounted onto glass slides for immunohistochemistry analysis.

Sections were deparaffinized in xylene and rehydrated in a descending ethanol series prior to heat-induced antigen unmasking in Citrate Buffer pH 6.0 (Sigma). After blocking of nonspecific binding in PBS-T containing 10% goat serum and 1% BSA, the samples were incubated with primary antibodies against TP53 (Santa Cruz Biotechnology, sc-126, 1:50), SARS-CoV-2 spike glycoprotein (Abcam, ab272504, 1:500), or CXCL8 (CST, 94407, 1:200) diluted in PBS-T containing 1% BSA overnight at 4°C in a humidified staining tray overnight. The next day, samples were incubated with secondary peroxidase-conjugated antibody (SignalStain Boost IHC Detection Reagent, HRP Mouse [8125S] or Rabbit [8114S], CST). Chromogenic detection was performed using the SignalStain DAB Substrate Kit (CST #8059) followed by hematoxylin counterstaining and dehydration in an ascending ethanol row. Slides were mounted with Eukitt Quick-hardening mounting medium (Sigma). Sections were imaged at 20x magnification using an Olympus BX-UCB whole slide scanner equipped with Olympus UPLSAPO and VS-ASW v2.7 software, and representative images were shown using OMERO^149^.

Immunohistochemistry quantification of TP53 was performed using QuPath 0.3.0. A patient lung tissue image was randomly selected to train an algorithm to detect cells and DAB staining. DAB intensities were Z-scaled across all patient samples. Cells with Z-scaled DAB intensities > 15 were considered TP53 positive. The percentage of TP53 positive cells for each patient was calculated by dividing the total number of positive nuclei by the total number of nuclei detected on the image and multiplying by 100.

### Cell fusion assay

The NanoBiT luciferase complementation system was used to quantify cell-cell fusion. To each well of a 96-well plate, 5e3 VeroE6 pLenti-HiBiT-BSD and 5e3 VeroE6 pLenti-LgBiT-P2A-BSD cells were added. Cell mixtures were reverse transfected with pcDNA3.1 expression plasmids via TransIT-X2 (Mirus) at a 1:3 ratio (50 ng plasmid, 0.15 μL TransIT-X2, and 5 μL OptiMEM per well). After 48 hr, media was removed, and 100 μL NanoGlo substrate (Promega) in OptiMEM (Thermo) was added to each well. Luminescence was measured with a SpectraMax iD3 or Glomax plate reader.

### Cell proliferation assay

In brief, 5e3 A549, A549ACE2, or A549ACE2 TP53ko pInducer20 cells were plated in each well of a 96-well plate. One day later, doxycycline-containing media was added at a final concentration of 1 μg/mL. After 72 hr, media was removed, and a 1:1 mixture of CellTiter-Glo reagent (Promega) and PBS was added to each well, incubated for 10 min at room temperature, and assayed for luminescence with a SpectraMax iD3 plate reader.

### RT-qPCR

A total of 300 ng RNA was used for cDNA synthesis using the SuperScript IV Reverse Transcriptase kit (ThermoFisher Scientific; #18090050). qPCR experiments were performed in a 384-well plate using LightCycler 480 SYBR Green (Roche; #4887352001) in a Roche LightCycler 480 II PCR system utilizing a Roche LightCycler 480 Software v1.5.1.62. QuantiTect Primer assays [*ACTB* (QT00095431) and *CDKN1A* (QT00062090)] we purchased from Qiagen. *ACTB* was used for normalization, and data were analyzed via the 2^-ΔΔCt^ method.

### Senescence-associated beta-galactosidase assay

The fluorescence-based senescence beta-galactosidase activity assay (CST #23833) was used to determine beta-galactosidase activity according to the manufacturer’s guidelines. In brief, 2.5e4 A549, A549ACE2, or A549ACE2 TP53ko pInducer20 cells were plated in each well of a 24-well plate. After 24 hr, media was swapped, and doxycycline was added at a final concentration of 1 μg/mL. After 48 hr, cells were washed with 1xPBS and lysed with senescence cell lysis buffer (CST #91029) containing 1x protease/phosphatase inhibitors (Thermo). After clarifying the supernatant, 50 μL 2x senescence reaction buffer (CST #78494) containing SA-beta-Gal substrate (CST #45954) and 10 mM beta-mercaptoethanol was added to 50 μL lysate and incubated at 37°C for two hours. 50 μL of the reaction mix was then added to 50 μL 2x senescence stop solution, and fluorescence was read with excitation at 360 nm and emission at 465 nm with a SpectraMax iD3 plate reader. In parallel, lysates were quantified by BCA assay (Thermo), and protein concentration was used to normalize senescence-associated beta-galactosidase activity.

### Small RNA sequencing

A total of 100 ng total RNA was used for library preparation via the QIAseq miRNA Library Kit (#331502, Qiagen) according to manufacturer’s recommendations. Briefly, QIAseq miRNA 3’ and 5’ adapters were ligated to mature miRNAs. The ligated miRNAs were then reverse transcribed to cDNA using a reverse transcription (RT) primer with a Unique Molecular Index (UMI). Libraries underwent cleanup and amplification (16 PCR cycles) using a forward primer that has a unique index and the reverse primer has another unique index for dual indexing (QIAseq miRNA 96 Index Kit IL UDI, #331645, Qiagen). Successful library production, quality control, and quantification was performed using Tapestation (High Sensitivity D1000 ScreenTape). Libraries were pooled together and subjected to sequencing via a NovaSeq 6000.

Raw sequencing reads were aligned to mature microRNA human sequences from miRbase v22.1 using bowtie v1.2.2 using options “-n 0 -l 30 --norc --best --strata -m 1”. UMIs were extracted with fgbio v2.1.0 function “AnnotateBamWithUmis” and duplicates removed with picard v2.8.0 MarkDuplicates. Reads were determined by samtools v1.3.1. Differential expression analysis was performed using DESeq2 in R.

### Cytokine profiling assay

In brief, 1e5 A549ACE2 or A549ACE2 TP53ko pInducer20 cells were plated in each well of a 6-well plate. After 24 hr, media was swapped with doxycycline-containing media at a final concentration of 1 μg/mL. After 48 hr, media was clarified for cytokine analysis via the Proteome Profiler Human Cytokine Array Kit (R&D Systems) as per the manufacturer’s instructions. Membranes were imaged on an Amersham Imager 600 (GE Healthcare), and spots were quantified using Fiji/ImageJ v2.3.0.

### Statistical analysis

No statistical methods were used to predetermine sample size. The experiments were not randomized. The investigators were not blinded to allocation during experiments and outcome assessment. All statistical analyses were performed in R^150^. Two-tailed statistical tests were used unless stated otherwise. Multiple comparison adjustments were performed as noted. Replicates were taken from distinct samples.

